# Evolutionary history of the main extracellular matrix polysaccharides in brown algae

**DOI:** 10.1101/2024.04.23.590721

**Authors:** Lisa Hervé, Ahlem Bouguerba-Collin, J. Mark Cock, France Denoeud, Olivier Godfroy, Loraine Brillet-Guéguen, Tristan Barbeyron, Agnieszka P. Lipinska, Ludovic Delage, Erwan Corre, Elodie Drula, Bernard Henrissat, Mirjam Czjzek, Nicolas Terrapon, Cécile Hervé

## Abstract

Brown algae belong to the Stramenopiles phylum and are phylogenetically distant from plants and other multicellular organisms. This independent evolutionary history has shaped brown algae with numerous metabolic characteristics specific to this group, including the synthesis of peculiar polysaccharides contained in their extracellular matrix (ECM). Alginates and fucose-containing sulphated polysaccharides (FCSP), the latter including fucans, are the main components of ECMs. However, the metabolic pathways of these polysaccharides remain poorly described due to a lack of genomic data. An extensive genomic dataset has been recently released for brown algae and their close sister species. We performed an expert annotation of key genes involved in ECM-carbohydrate metabolisms, combined with comparative genomics, phylogenetics analyses, and protein modelling. Our analysis indicates that the gene families involved in both the synthesis and degradation of alginate were acquired by the common ancestor of brown algae and their closest sister species *Schizocladia ischiensis*, and subsequently expanded in brown algae. The pathway for the biosynthesis of fucans still remains biochemically unresolved and we identify the most likely fucosyltransferase genes that may harbour a fucan synthase activity in brown algae. Our analysis questions the possible occurrence of FCSPs outside brown algae, notably within their closest sister taxon.

## Introduction

Brown algae correspond to the class Phaeophyceae within the Stramenopile lineage (1). Many of these organisms are key components of extensive coastal ecosystems that provide high value ecosystem services, including the sequestration of several megatons of carbon per year. They represent a rich but underutilized biomass that will likely provide solutions to address global challenges, such as climate change, food shortages, and rising demands for natural bioactive compounds (2,3). These organisms are also impacted by climate change and shifts in their habitats and communities have been reported (4,5). They are key components of highly dynamic ecosystems, and to thrive in such habitats they have evolved plasticity in terms of metabolic responses, including at the level of their extracellular matrices (ECMs), or cell walls (6).

Brown algae evolved complex multicellularity independently from animals, fungi and land plants, and their metabolic networks have been shaped by secondary endosymbiosis and a number of horizontal gene transfers (HGTs) (6–8). This has resulted in the emergence of a range of unique metabolic features, including the generation of specific glycans in their ECMs (9,10). Alginates and fucose-containing sulfated polysaccharides (FCSPs) are the main components of the ECM, representing up to 45% of algal dry weight. Alginates are linear polymers made solely of two 1,4-linked epimers: β-D-mannuronic acid (M) and α-L-guluronic acid (G). FCSPs include both fucans and fucoidans (9). Fucans are highly sulfated polysaccharides with a backbone structure based on sulfated L-fucose residues, on which additional branches of sulfated fucose, galactose and glucuronate can occur. Fucoidans are a set of heterogeneous polymers with non-fucose backbones (e.g. chains of galactose, mannose or glucuronate), and branches of sulfated fucose (9).

The metabolic pathways for both alginates and sulfated fucans have been predicted (11,12). All enzymes biochemically characterized so far are involved either in the synthesis of sugar precursors or in the remodelling of alginates (12). GDP-mannose is an essential activated sugar used in the alginate biosynthetic pathway, and in the production of fucans through its conversion to GDP-fucose (12). The three enzymatic steps that lead to the production of GDP-mannose from fructose-6 phosphate are catalysed by mannose-6-phosphate isomerase (MPI), bifunctional phosphomannomutase (PMM)/phosphoglucomutase (PGM) and mannose-1-phosphate guyanylyltransferase (MPG). The later enzyme also has MPI activity (12,13). As in other eukaryotes and in most bacteria, the precursor GDP-fucose is likely produced in brown algae from GDP-mannose (the *de novo* pathway) and from fucose (the salvage pathway) (12). In the *de novo* pathway, GDP-mannose is first converted into GDP-4-keto-6-deoxymannose by GDP-mannose 4,6-dehydratase (GM46D), and subsequently converted to GDP-fucose by GDP-fucose synthase (GFS). In the salvage pathway, cytosolic fucose is first phosphorylated by a fucokinase (FK) and then condensed to GTP by a GDP-fucose pyrophosphorylase (GFPP) to form GDP-fucose. Once GDP-mannose has been synthesized, its oxidation to GDP-mannuronic acid will result in the elongation of an alginate polymer in the form of polymannuronate. Oxidation of GDP-mannose is catalyzed by GDP-mannose dehydrogenase (GMD).

Once these activated sugars (e.g. GDP-mannose, GDP-fucose) have been generated, the elongation or grafting of the glucan chains is the most significant step in ECM synthesis, but, for brown algae, the genes involved have not been identified (12). Carbohydrate active enzymes (CAZymes) are major actors in polysaccharide metabolism (14,15). They are classified into different families in the CAZy database (www.cazy.org, (14)) based on their amino acid sequence similarities. CAZy classification includes glycosyltransferases (GTs), glycoside hydrolases (GHs), polysaccharide lyases (PLs), carbohydrate esterases (CEs), enzymes with Auxiliary Activities (AAs) and associated modules such as Carbohydrate-Binding Modules (CBMs). Additional important enzymes involved in the remodeling of glycans in brown algae include mannuronan C5-epimerases which convert M to G residues in alginates, and the sulfotransferases (STs) and sulfatases (SAs), which mediate the sulfation and desulfation of FCSPs, respectively (7,12). The SAs have been classified into families in the SulfAtlas database based on amino acid sequence similarities (https://sulfatlas.sb-roscoff.fr, (16,17)).

Sequencing a large number of genomes of brown algae and their sister groups, allows the emergence and diversification of key biological traits within the brown algae to be investigated, particularly traits associated with the transition to complex multicellularity, such as the elaboration of an adherent ECM. Brown seaweeds have remained poorly described in terms of genome sequencing due, in part, to difficulties with extracting nucleic acids. The Phaeoexplorer project has recently generated 60 new genomes corresponding to 40 species of brown algae and four close sister species (https://phaeoexplorer.sb-roscoff.fr, (18)). The 40 brown algal species include representatives of 16 families, spanning all the major orders of the Phaeophyceae. The sister species notably include the sister taxon *Schizocladia ischiensis*, the closest taxon to the brown algae (1,18,19). Here, we have investigated the evolutionary history of carbohydrate metabolism gene families in brown algae by a combination of expert annotations and evolutionary analysis of CAZymes (14), STs and SAs (16), as well as protein modelling of the most promising candidate genes. When relevant, we also performed genomic analyses of additional eukaryotes such as echinoderms or diatoms. This study offers the first representative view of the core CAZome in brown algae, derived from an extensive resource of sequences from distinct taxonomic origins. Our analysis indicates that the abilities to both synthesize and degrade alginate were gained prior to the diversification of the brown algae. We also identify the most likely fucan synthase genes in brown algae, an enzymatic step that still remains biochemically unresolved.

## Material and methods

### Identification of carbohydrate-related proteins

The CAZyme predicted proteins were identified by human curators of the CAZy database (http://www.cazy.org/, (14)). Curation consisted in the expert validation of the predicted modularity for each protein, using a combination of BLASTp pairwise alignments against both characterized members and previously curated sequences integrated in the CAZy database, as well as HMMER v3.4 (www.hmmer.org; (20)) comparisons to in-house HMM profiles of the families, and subfamilies when available. Proteins obtaining 100% coverage without gaps in pairwise alignments against a previously curated sequence were not manually inspected, allowing to decrease the load of human curation by processing genomes sequentially. The ST predicted proteins from various Stramenopiles were identified in the UniProt database based on the presence of a sulfotransferase-specific Pfam signature. Sequences of brown algal genomes not present in Uniprot were obtained by BLASTp searches in the Phaeoexplorer database (https://phaeoexplorer.sb-roscoff.fr, (18)) using the sequences found in the UniProt database as query. All other genes were identified by BLASTp searches of brown algal enzymes in the Phaeoexplorer database, using characterized brown algal genes as the queries, when known (MPI, PMM, GMD, UGD, ManC5-E, DEHU reductase). The homology threshold was chosen as ∼35% pairwise sequence identity. In addition to the identified brown algal sequences, representative and characterized members of the targeted families were retrieved from the CAZy database (for CAZymes) and from public databases.

### Phylogenetic analyses

Protein sequences were aligned with the MAFFT program using the FFT-NS-i iterative refinement method (https://mafft.cbrc.jp/alignment/server, (21)). In the case of multimodular proteins, the targeted catalytic domains were extracted prior to their alignment. The multiple alignments were curated manually using the Jalview program (https://www.jalview.org, (22)) to retain only the informative sites. Prior to carrying out comprehensive phylogenetics, a preliminary analysis was carried out for each protein family to select the optimal model of sequence evolution using the IQ-TREE progam (http://iqtree.cibiv.univie.ac.at/; (23)). Phylogenetic trees were subsequently constructed using the maximum likelihood approach with the RAxML program ((24), available at https://usegalaxy.fr). The reliability of the trees was tested with an MR-based Bootstopping criterion which generated resamplings of the datasets (range of 350 to 1000). The trees were edited in Mega 11.0.13 (https://www.megasoftware.net/; (25)) and graphically shaped with Adobe Illustrator CS6. For all phylogenies, only bootstrap values above 60% are shown. The brown algal sequences are indicated with a color code in relation to their taxonomy as indicated by the key legend. Sequences from the closest sister taxa *S. ischiensis* are further indicated by the red circle. In all instances, the green squares indicate sequences which have been biochemically characterized.

### Heatmaps based on Blast Score Ratios

To calculate the Blast Score Ratio (BSR), all pairs of sequences were compared using the NCBI BLASTp tool. The BSR, which corresponds to the bitscore normalized to the length of the protein query, was then calculated from the BLAST result following the procedure described in (26). Heat maps representing these BSR results and associated dendrograms were constructed using the ggheatmap function in the ggplot2 package in RStudio.

### Extracted alignments and AlphaFold2 structural modeling

Some specific catalytic regions were extracted from alignments (Figures 2C, 5B, 8C). Alignments of full-length proteins were made using the MAFFT program and edited in the ESPript 3.0 program (https://espript.ibcp.fr; (27)). The color code of the amino acid residues indicates their biochemical character. Regions displaying high sequence similarity are boxed. The green squares indicate sequences which have been biochemically characterized. AlphaFold v2 (28) was used to predict 3D protein structures from primary amino acid sequences. The PyMOLv2.5.5 Molecular Graphics program (http://www.pymol.org/) was used to create and the display structural images.

### Syntenic region analysis

Orthogroups, derived from predicted proteins of the 60 strains sequences in the Phaeoexplorer project (https://phaeoexplorer.sb-roscoff.fr, (18)) were searched for genes of interest. Microsynteny analysis was carried out on a subset of species where the target orthologues resided on a singular chromosome/scaffold, using McscanX (29), following the pipeline described at https://github.com/tanghaibao/jcvi/wiki/Mcscan. Gene annotations were then processed using AGAT (https://github.com/NBISweden/AGAT) and LASTZ (30) tools (C-score cutoff =0.7 to infer synteny blocks between species.

## Results

### CAZyme profiles in brown algae

Expert annotation of carbohydrate metabolism gene families was performed for all the major orders of brown algae (Phaeophyceae) and their close sister groups. Regarding CAZymes, we have already reported a quite constant number of genes (237 genes on average) for the 7 orders of brown algae (Figure 1A) (18). This number is similar in the closest sister taxon *S. ischiensis* (233 genes) while it is twice more than the more distantly related unicellular alga *Heterosigma akashiwo* (120 genes). The brown algal and sister taxa CAZymes are distributed across 69 families (including 37 GT, 21 GH and 1 PL families) with, on average, three genes per family (while reaching up to 16 genes for GT2 and GT4 and 19 genes for the PL41) (Figure 1B). The GT31 and GT47 families which are abundant in Viridiplantae (essentially β-galactosyl- or β-glucuronyltransferases, (31,32)) are also among the largest GT families in brown algae (average of 9 genes per genome for both families). We also detected in brown algae the occurrence of unexpected CAZyme families such as GT23 (α-1,6-L-fucosyltransferase, (33,34)), usually found in metazoan, GT49 (α-glucuronyltransferase), found in sac fungi, GT60 (α-N-acetyl-glucosaminyltransferase), found in amoeba plus chlorophyte green algae. Regarding glycosyl hydrolases, the most populated families were GH16, GH47, GH81 and GH114 (with 6, 5, 4 and 5 genes per genome respectively). The GH16 and GH81 enzymes are likely involved in the metabolism of the storage polysaccharide laminarin in brown algae, with β-1,3-glucanase activities already referenced in those families. GH47 genes are frequently found in eukaryotic genomes and are associated with α-1,2-mannosidase activity. Outside brown algae, GH114 are found in chlorophytes, fungi and bacteria, with a reported α-1,4-galactosaminidase activity (35,36). Finally PL41, which is, so far, the only PL family reported in brown algae and with alginate lyase activity (37), contains a high number of genes.

**Figure 1.**
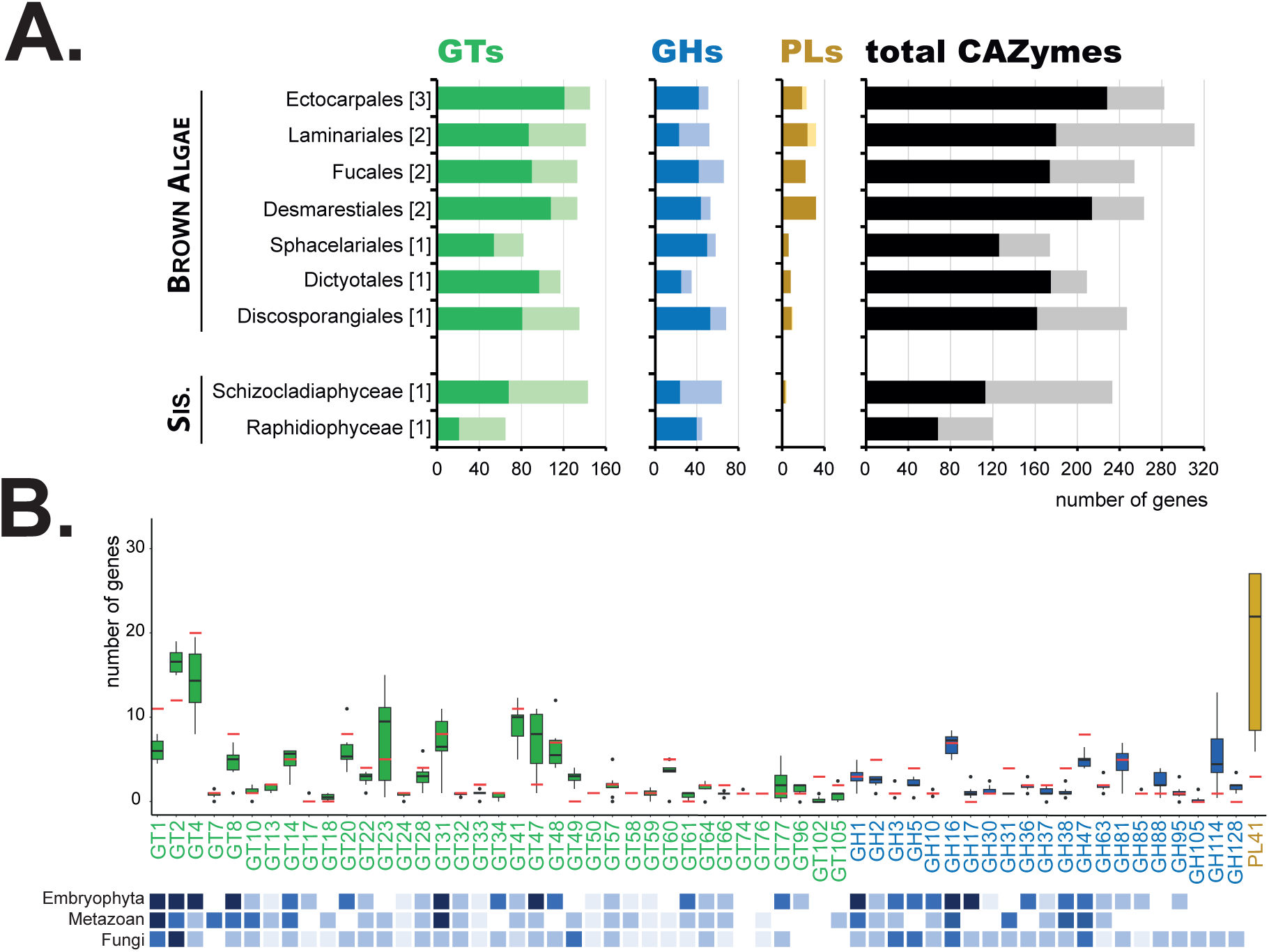
Overview of CAZyme families in brown algae. (**A**) Counts of numbers of genes predicted to encode glycosyltransferases (GTs), glycoside hydrolases (GHs), polysaccharide lyases (PLs) and for all CAZymes (GTs, GHs, PLs, Carbohydrate Esterases CEs, Auxiliary Activities AAs, Carbohydrate Binding Modules CBMs), showing numbers of both full-length proteins (dark colours) and fragments (light colours). The data is averaged by taxonomic order with the number of species analysed per order in brackets. (**B**) Number of genes for selected CAZyme families in brown algae (box plots) and the sister species *S. ischiensis* (red lines). Counts include non-fragmentary proteins and fragments. The species analysed are the same as in Supplementary Table S1. A schematic overview regarding abundance of the corresponding families is indicated below the plots for other groups (the reader is directed to the CAZy database for a more accurate counting in these other groups).

### The genes involved in alginate metabolism in brown algae are also present in its closest sister taxon *S. ischiensis*

The alginate and the *de novo* fucan biosynthetic pathways share common steps leading to the generation of GDP-mannose as a precursor, prior to its conversion to GDP-fucose in the case of fucans. The two enzymes involved in these enzymatic conversions, MPI/MPG and PMM/PGM, have been characterized for *Saccharina japonica* (13,38). Analysis of their phylogeny indicated that these enzymes are widely present in brown algae but also more generally within the Stramenopiles, with several occurrences in the closest sister taxa and diatoms (Supplementary Figure S1A,B). GDP-mannose dehydrogenase (GMD) is essential to provide GDP-mannuronic acid for the initial assembly of alginate in its mannuronan form by a mannuronan synthase. The latter putatively belongs to the large and multispecific GT2 family, although no functional validation has been obtained so far to identify specific candidate genes, making phylogenetic studies of this family highly speculative. The alginate obtained undergoes further modelling through the action of mannuronan C5-epimerases (ManC5-E). Both GMD and ManC5-E activities have been described, including in brown algae (13,39–42).

GMD belong to a protein superfamily also including and uridine diphosphate-glucose dehydrogenases (UGD). In order to explore the sequence diversity within this family, we gathered GMD and UGD protein sequences, using representative sets of brown algal sequences and additional characterized members from public databases. After clustering their BLASTp+ Score Ratio (BSR), we generated heatmaps that clearly illustrates that UGD and GMD form two distinct clusters (Supplementary Figure S2A). Within the first cluster, the UGD sequences from brown algae and the sister taxon *S. ischiensis*, share more than 90% pairwise protein sequence identity. In the second cluster, GMD sequences from brown algae and the sister taxon *S. ischiensis*, share more than 77% pairwise protein sequence identity, while they share less than 25% identity with the UGD sequences, indicating a large evolutive distance between these two families. The BSR-based heatmap indicates a distant cluster of sequences from *Actinobacteria*, which share around 30% identity with both GMD and UGD brown algal sequences. An additional BLAST search indicated that the brown algal GMDs share stronger sequence identities (40%) with a protein belonging to *Candidatus woesebacteriota* (MBP7967557.1), an uncultivated bacterium from a putative novel taxon (43). The phylogeny suggests that the common ancestor of *S. ischiensis* and brown algae probably gained the GMD sequence via horizontal gene transfer (HGT), and that the gene was duplicated in all brown algal groups (Figure 2A, Supplementary Figure S2B). The mannuronan C5 epimerases are a large multigenic family in brown algae, with an average of 27 members identified per genome. Eleven homologues were identified in *S. ischiensis* (Figure 2B, Supplementary Figure S3). Unlike aforementioned families, screening the other Stramenopile genomes available (*i.e*. other brown algal sister taxa, diatoms and oomycetes), did not identify any ManC5-E sequences. Three ManC5-E orthologue groups were identified (Figure 2B, Supplementary Figure S3). All the ManC5-E sequences possess the DPHD motif, typical of the ManC5-E catalytic site (Figure 2C), indicating that these sequences are genuine ManC5-Es. Marked amplifications of the gene family were detected in the Fucales, Desmarestiales and Laminariales. The closest bacterial representatives are from *Actinobacteria* with ∼30% protein pairwise identity (Figure 2B, Supplementary Figure S3).

**Figure 2.**
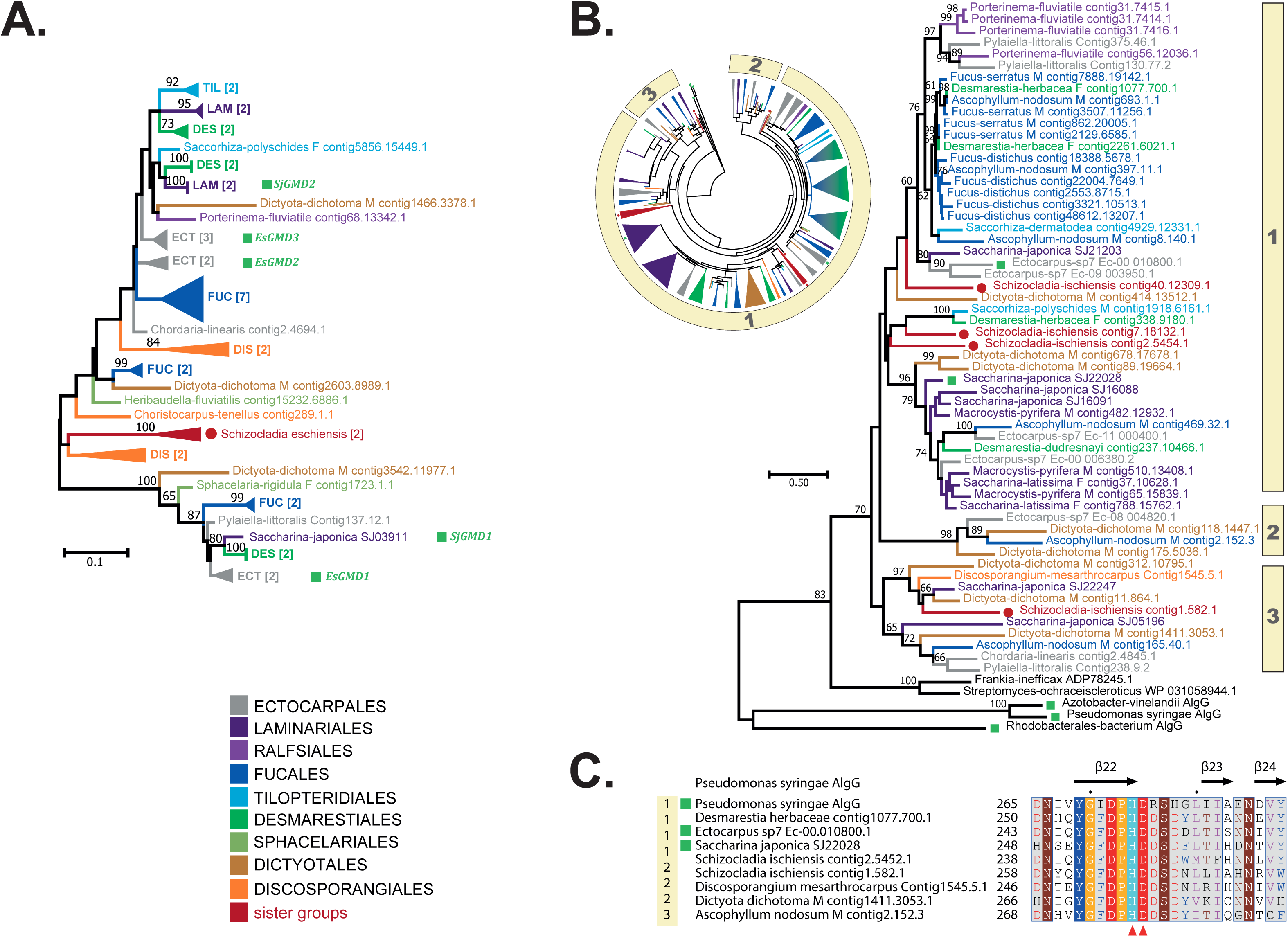
Phylogenetic trees of key enzymes involved in the synthesis of alginates. (**A**) Phylogenetic tree of GDP-mannose 6-dehydrogenases (GMD). The uncompressed tree is shown in Supplementary Figure S2B. (**B**) Phylogenetic trees of mannuronan C5-epimerases (ManC5-E). The circle shown as an inset indicates a global phylogeny of a large collection of ManC5-Es from various genomes, in which 3 main orthologous groups can be found. The phylogeny shown on the right side is representative of this global view, the 3 groups being indicated. The uncompressed tree of the inset is shown in Supplementary Figure S3. (**C**) Extract of an alignment of selected ManC5-E sequences from brown algae and the sister species *S. ischiensis*, the bacterial ManC5-E sequence from *Pseudomonas syringae* (pdbcode: 4NK6). The region shown corresponds to the catalytic site as identified in 4NK6, and the secondary structure assignments shown above the sequences are those determined for 4NK6. The red arrowheads below the alignment indicate the conserved catalytic residues His274 and Asp275. The numbering on the left refers to the phylogenetic clusters indicated in (B).

In addition to alginate synthesis, brown algae also possess a metabolic pathway for alginate degradation, the main actors being the PL41s. This is one of the largest degrading CAZyme family in brown algae (Figure 1B), with an average of 19 genes per genome, and can be classified into at least four orthologous groups. PL41 duplication events have occurred in several brown algal orders including the Ectocarpales and the Laminariales. *S. ischiensis* displays three PL41 sequences while no homologues were found in other Stramenopiles (Figure 3A, Supplementary Figure S4A). PL41s are however not limited to brown algae, with bacterial members in some *Gammaproteobacteria*, *Actinobacteria* and *Alphaproteobacteria*, but with an average of one gene copy per genome (34-40% pairwise identify with the PL41 domain of brown algae). The oligoalginates generated by PL41s have been shown to be further degraded into 2-keto-3-deoxy-gluconate (KGD) by a DEHU reductase in the brown alga *S. japonica* (44). DEHU homologues were found in most brown algal genomes but not all sister taxa. This family of genes is found in other eukaryote genomes (Supplementary Figure S5). Overall, the broad distribution of this family of enzymes across eukaryotic organisms and the lack of homologues in some brown algae, argue against a strong specificity for alginate degradation in brown algae.

**Figure 3.**
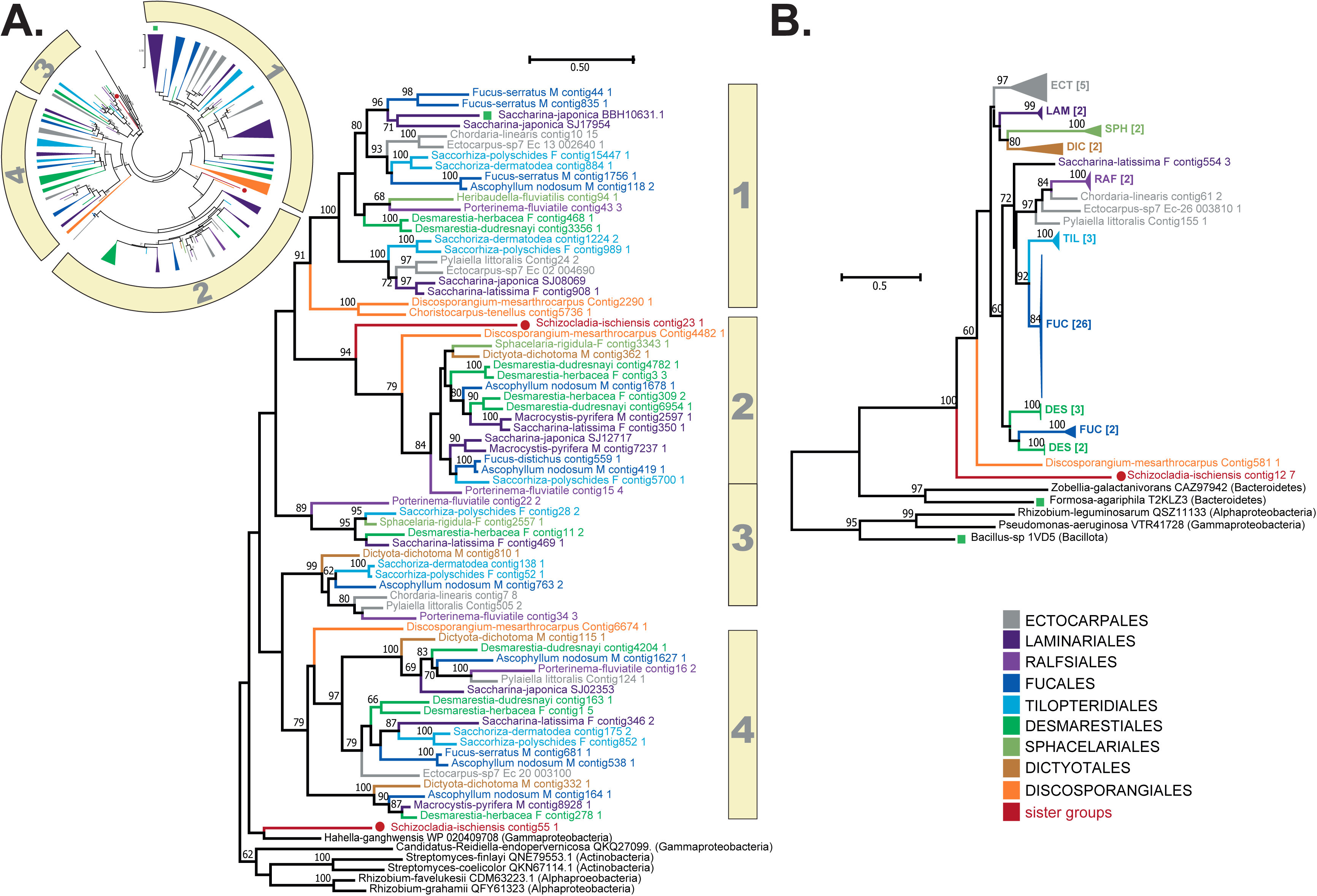
Phylogenetic trees of key enzymes involved in the degradation of alginates. (**A**) Phylogenetic tree of alginate lyases from the PL41 family. The circle shown as an inset indicates a global phylogeny of the large collection of PL41s from various genomes, in which four main orthologue groups can be found. The phylogeny shown on the right is representative of this global view, the four groups being indicated. The uncompressed tree of the inset is shown in Supplementary Figure S4. (**B**) Phylogenetic tree of glycoside hydrolases from the GH88 family. The uncompressed trees are shown in Supplementary Figure S6.

GH88 family members were found in all the brown algal genomes analysed (Figure 3B, Supplementary Figure S6). This CAZyme family includes characterized members which specifically act on oligosaccharides with an unsaturated uronic acid moiety (Δ), typically introduced by the lytic mechanism of lyases, such as heparan lyases (45) or ulvan lyases (46). As the PL41 alginate lyases are the only known PL family described so far in brown algae, the GH88 family would be an ideal candidate for further hydrolysis of oligosaccharides generated by PL41 action. The GH88 family in brown algae shares a common ancestor with that of its sister taxon *S. ischiensis* and the family has amplified in brown algae (Figure 3B, Supplementary Figure S6), especially within the Fucales where it is a large family (around 16 genes per genome).

### Prediction of candidate genes for the synthesis of sulfated fucans

All the candidate genes putatively involved in the synthesis of GDP-fucose await functional validation in brown algae. Genes encoding the *de novo* pathway enzymes GM46D and GFS were identified in all the orders of brown algae analysed, with close homologues in sister taxa, including *S. ischiensis* (Supplementary Figure S7A,B). In contrast, genes encoding fucokinase (FK) and GDP-fucose pyrophophorylase (GFPP) from the salvage pathway were identified in the Phaeophyceae, but not in *S. ischiensis* (Supplementary Figure S7C,D). These two enzymes are usually found as a bimodular protein (FK-GFPP) in brown algae. They are found in a gene cluster previously reported in three Ectocarpales, which also encodes a sulfotransferase (ST) and a hydrolase (47). This microsyntenic region was found in other members of the Ectocarpales (Figure 4) and is conserved in other brown algal orders (Laminariales, Fucales, Desmarestiales), albeit with a stronger distance between the contained genes near the root (*Dictyota dichotoma*). The synteny was not observed with the Discosporangiales and more distant taxa (sister taxa).

**Figure 4.**
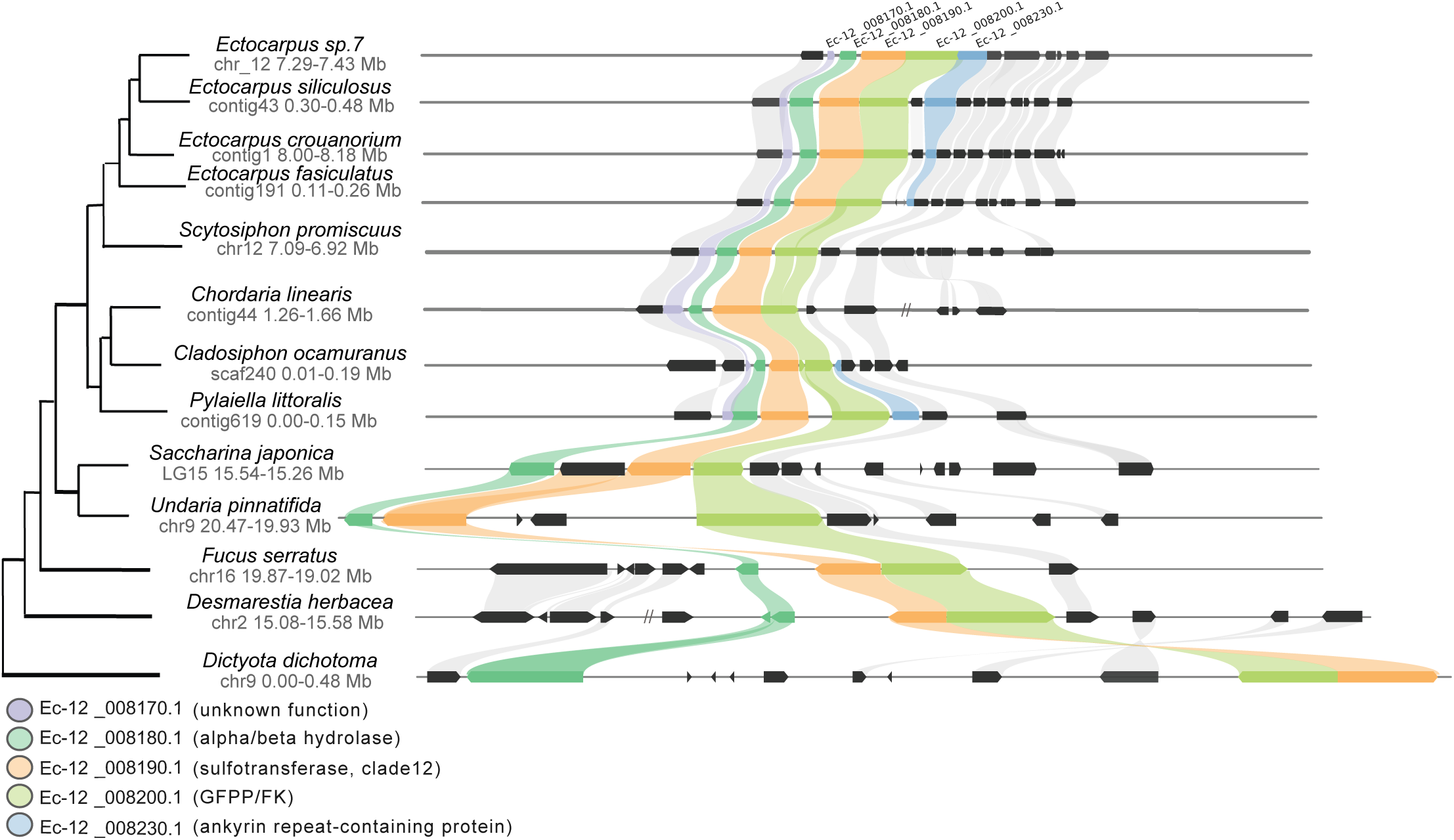
Syntenic region of putative fucan-related genes in brown algal genomes. This region contains five conserved genes encoding: an Ankyrin repeat-containing protein, a GDP-fucose pyrophosphorylase (GFPP), a L-fucokinase (FK), a sulfotransferase (ST), an alpha/beta hydrolase.

The most important set of enzymes involved in the synthesis of fucans are the fucosyltransferases (FucTs), which transfer GDP-fucose to oligofucans to elongate the chains. Their exact nature is still unknown. The CAZy database includes several FucTs that have been identified and these are classified into a variety of GT families. Our previous analyses of a restricted number of brown algal genomes, indicated that our FucT candidate genes for fucan synthesis may belong to either the GT10, GT23, GT41 and GT74 families (12). The extensive Phaeoexplorer dataset allowed us to performed a more in-depth analyses of these CAZyme families in brown algae, and to refine the identification of putative fucan synthases (Supplementary Table S1).

GT10s are referenced as α-1,3 or α-1,4-FucTs in the CAZy database, and the activities described relate to the transfer of an N-acetyl glucosamine (GlcNAc) residue (48,49). The fucose residues in fucans are α-1,3 and/or α-1,4-linked. It is thus tempting to hypothesise that the GT10 family is involved in their synthesis. However, there is only one GT10 gene per brown algal genome on average (Figure 5A, Supplementary Figure S8, Supplementary Table S1). Considering the importance of sulfated fucans in ECMs of all brown algae, we would expect the corresponding FucT gene to be present in multiple copies. As such this GT10 family is not the most favorable candidate for their biosynthesis. Moreover, a close homologue of brown algal GT10s is found in the Pelagophyceae, which are known not to contain fucans. The phylogeny further indicates that the brown algal sequences are closely related to plant GT10s, which are involved in protein glycosylation, where the α-1-3 FucT activity takes place on a GlcNAc residue, directly linked to an asparagine (50). While the activity of brown algal GT10s on oligofucans is unlikely, the GDP-fucose binding site is nonetheless highly conserved (Figure 5B,C), indicative of FucT activity. Comparisons were made with the only known structure of a GT10 from bacteria (51), and additional animal GT10 sequences for which the donor binding site has been studied (52,53). Both the alignment (Figure 5B) and the structure predictions (Figure 5C) indicate the conservation of key residues within the donor site in the brown algal sequences, while the acceptor site exhibits considerable variation (Figure 5D,E).

**Figure 5.**
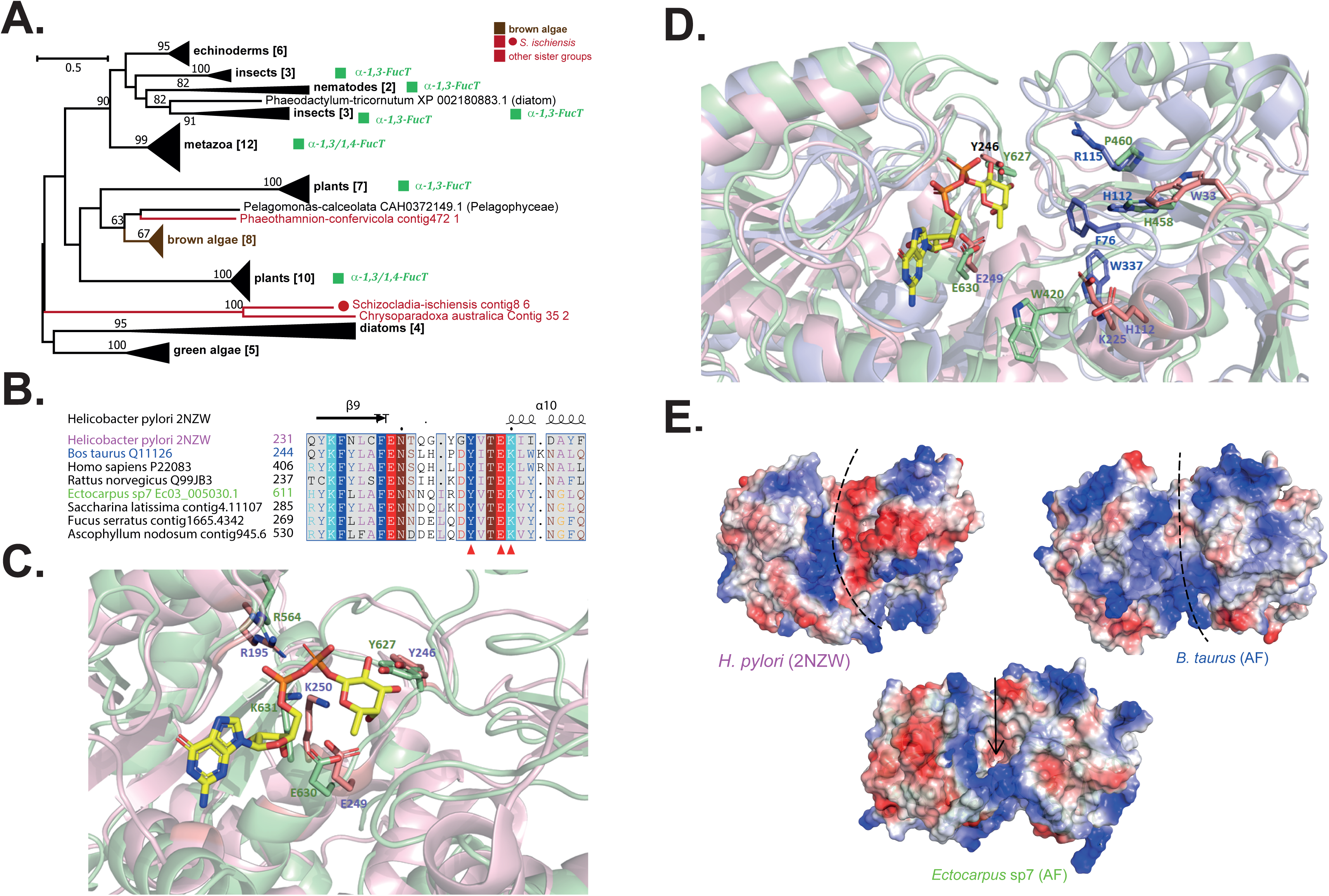
Comparison of brown algal GT10s with biochemically characterised orthologues. (A) Phylogenetic tree of the GT10 family. This family is known to contain FucT activities for species outside brown algae. The uncompressed tree is shown in Supplementary Figure S8. (B) Extract of an alignment of selected GT10 sequences from brown algae against the bacterial GT10 sequence from *Helicobacter pylori* (pdbcode: 2NZW) focusing on the GDP-fucose donor site. Additional animal sequences are also aligned and secondary structure information is shown for 2NZW. The red arrowheads below the alignment indicate the conserved catalytic residues Tyr246, Glu249 and Lys250, which interact with the donor substrate GDP-fucose. The 2NZW sequence is shown in pink, a *Bos taurus* sequence in blue and the *Ectocarpus* species 7 sequence in green, and the same color code is used in C, D and E. (**C**) Overlay of the *H. pylori* GT10 crystallized structure (2NZW; pink) and the best models generated by AlphaFold2 for the *B. taurus* (blue) and brown algal (green) sequences. The cartoon representation focuses on the GDP-fucose binding site where GDP-fucose is apparent in yellow/orange colors. Some conserved catalytic residues, including those discussed in B), are highlighted. (**D**) Larger view of the overlay presented in B) which allows visualization of the acceptor binding site, in addition to the GDP-fucose binding site. Despite apparent structural conservation, the nature of the residues at the pocket surface differ drastically between the 3 sequences. (**E**) Molecular surfaces of 2NZW and the AlphaFold2 predictions for the *B. taurus* and algal sequences based on their electrostatic potentials. While the substrate binding cleft is clearly apparent in 2NZW and in the *B. taurus* sequence (dotted lines), it appears more enclosed in the brown algal sequence (arrow), although the N-terminal domain may form a flexible loop to allow accommodation of a large substrate.

GT41 is a multispecific family in the CAZy database, with N-acetylglucosaminyl-transferase and protein O-fucosyltransferase activity, the latter being restricted, so far, to the green lineage (54). Brown algal genomes harbor ten GT41 genes on average, with homologues found in all ochrophytes, usually with several copies per genome. The GT41 phylogeny indicates that brown algal GT41s are organized into numerous orthologous groups, with *S. ischiensis* frequently found at the base of these algal groups. None of these orthologues are closely related to plant O-FucT (Figure 6A, Supplementary Figure S9). A significant number of the orthologous groups are distant from the orthologous groups that contain the GlcNAc characterized activity, and could indicate new activities in the family. Overall, the phylogeny shows that the evolution of O-FucT activity has been restricted to the green lineage so far. Finally, the presence of multiple GT41 homologues in ochrophytes other than brown algae, does not favour the hypothesis that this class of enzymes contains fucan synthases.

**Figure 6.**
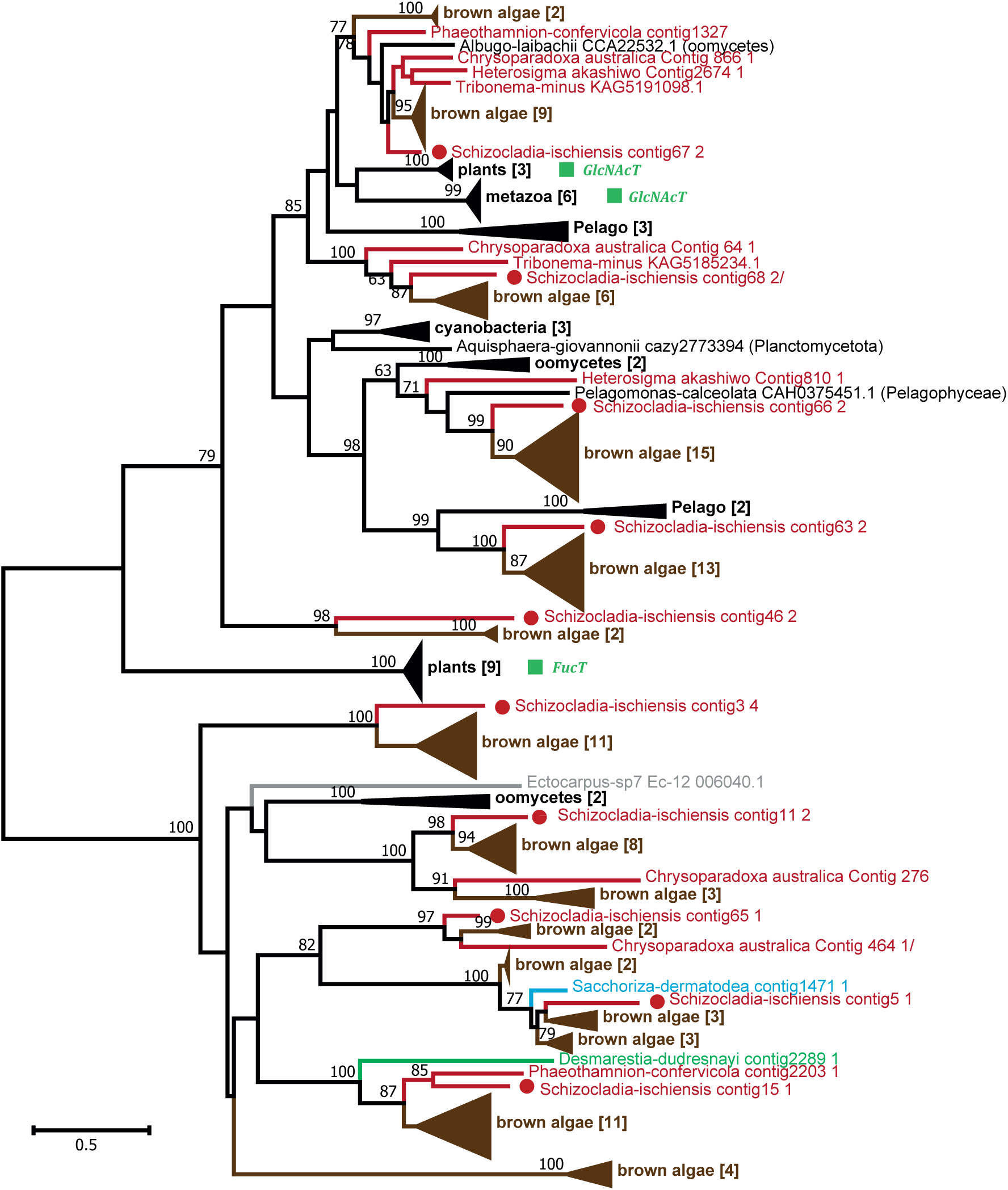
Phylogenetic tree of the GT41 family. Some GT41 family members in non-brown-algal species are known to have FucT activities (green boxes). The uncompressed tree is shown in Supplementary Figure S9.

α-1-2 fucosyltransferase activity has been reported in the GT74 family, which is also represented in brown algal genomes. This family has a restricted occurrence in eukaryotes, being only characterized in the amoeba *Dictyostelium discoideum*, where it participates in the glycosylation of the Skp-1 protein, involved in the ubiquitination regulatory system (55). The *D. discoideum* enzyme has a bimodular structure, the GT74 module being appended to a GT2 domain with β 1,3-galactosyltransferase activity. Bimodularity was also observed in the brown algal sequences, although the GT74 domain was found at the N-terminus, with the GT2 domain at the C-terminus, similarly to the situation found in some bacteria and the inverse of the situation in amoeba. Phylogenetic analysis indicated a common origin of GT74 genes from brown algae and other Stramenopiles (Figure 7, Supplementary Figure S10), although some genomes lacked this gene (*i.e.* some diatoms and brown algal sister taxa). Most brown algae have one GT74 gene. Again, considering the preponderance of sulfated fucans in ECMs of brown algae, and that this gene occurs in Stramenopiles known not to produce fucans, the GT74 family is not the best candidate to contain fucan synthases.

**Figure 7.**
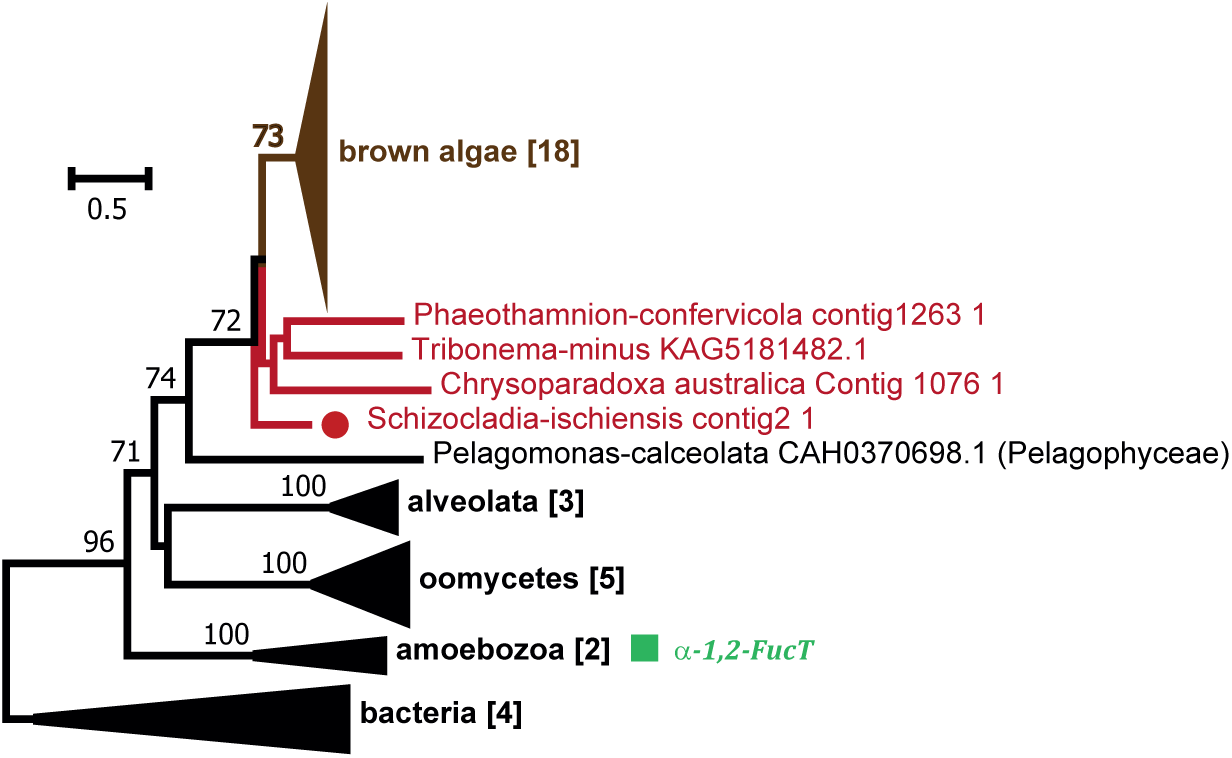
Phylogenetic tree of the GT74 family. Some GT74 family members in non-brown-algal species are known to have FucT activities (green box). These proteins frequently occur as bimodular forms with a GT2 domain. The phylogeny is based uniquely on the catalytic GT74 domain. The uncompressed tree is shown in Supplementary Figure S10.

Finally, the GT23 family is present in all brown algae and is known to contain fucosyltransferases in other eukaryotes. GT23 genes are usually found in metazoa, where α-1,6 FucT activity has been reported and the structure of the human isoform FUT8 has been characterised (33,34). Uncharacterized homologues have also been found in bacteria, fungi and the Chlorophyta but this family is very rare in these lineages. On average, seven GT23 genes were found per genome in brown algae (Supplementary Table S1). Characteristic GT23 features are conserved in the brown algal sequences but the proteins are only distantly related to other sequences in the family. A BSR-based heatmap indicated two main clusters (Figure 8A), with the brown algal sequences representing an independent cluster, sharing on average less than 30% pairwise identity with the second cluster that contains the metazoan sequences. The phylogeny of the brown algal GT23 sequences shows that they do not strictly distribute in relation to taxonomy and this may indicate the presence of orthologue groups (Figure 8B, Supplementary Figure S11). All these GT23 sequences share, on average, 37% pairwise protein identity. Some duplication events have occurred in the Fucales and Laminariales. Overall, this analysis indicated that brown algae contain GT23 sequences that may belong to one or to several sub-classes within the GT23 family. The diversity of brown algal GT23 proteins may indicate functionally and/or structurally distinct proteins. This diversity of proteins could potentially reflect the variety of positions to which a fucose is transferred in fucans and more generally in FCSPs (variety of monosaccharides, carbon position, linkage type). As such, the diverse GT23 proteins may reflect the FCSP diversity and the complex enzymatic machinery required for their synthesis.

**Figure 8.**
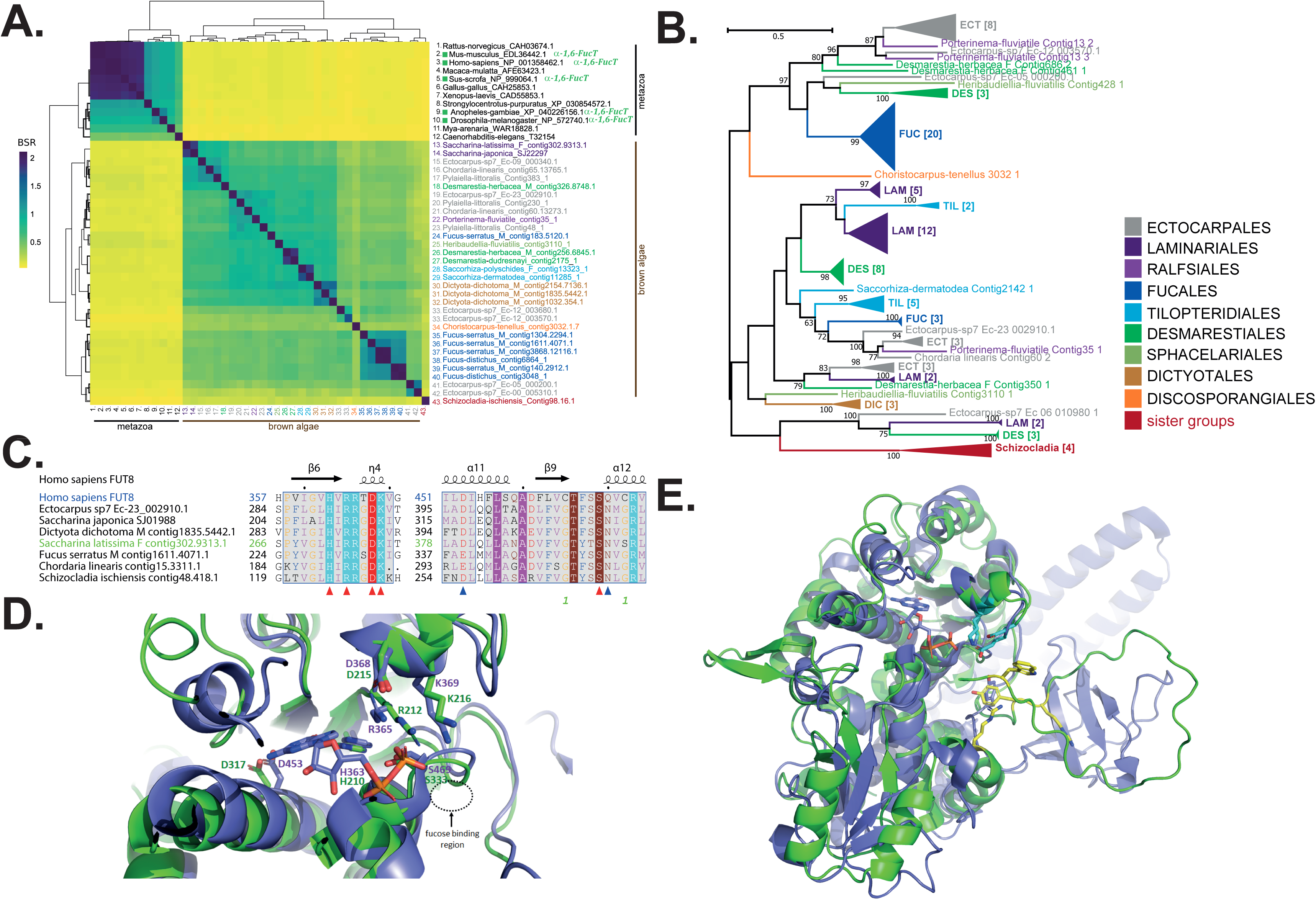
Comparison of brown algal GT23s with characterised orthologues. (**A**) Heatmap representing BLASTP+ Score Ratios (BSRs) for GT23 protein sequence pairwise alignments. The colour scale correlates BSRs. Two groups are apparent: one containing metazoan sequences and the other consisting of brown algal GT23s. (**B**) Phylogenetic tree of GT23s from brown algae and the sister taxa *S. ischiensis*. The uncompressed tree is shown in Supplementary Figure S11. (**C**) Extracts of an alignment of selected GT23 sequences from brown algae and the sister taxa *S. ischiensis*, plus the human FUT8 GT23 sequence (pdbcode: 6X5R), focusing on the GDP-fucose donor site as identified in FUT8. Secondary structure predictions for FUT8 are shown above the sequences. The red arrowheads below the alignment indicate the conserved catalytic residues His363, Arg365, Asp368, Lys369 and Ser469. The blue triangles indicate catalytic residues identified in 6X5R which differ in some of the algal sequences, but may retain similar biochemical properties: Asp453 and Gln470. The FUT8 sequence is shown in blue and the *S. latissima* in green, so as to match the color code used in D and E. (**D**) Overlay of the FUT8 crystal structure (6X5R) and the best model generated by AlphaFold2 of a brown algal sequence from *S. latissima* (Accession number: contig302.9313). The cartoon representation focuses on the GDP-fucose binding site where FUT8 is colored in blue and SlGT23A2D in green. The GDP is apparent in blue/orange colors. The conserved catalytic residues discussed in A) are indicated. (**E**) Global view of the overlay presented in D). The conserved catalytic residues Lys369 and E373 are shown in cyan. Aromatic residues present in the *S. latissima* sequence and possibly involved in acceptor fixation are shown in yellow.

As the brown algal GT23 sequences seem rather distant from other GT23s, and knowing that some of the latter have been biochemically and structurally characterized, we examined further the potential catalytic sites of the brown algal sequences. A sequence alignment was made for a selection of GT23 proteins from brown algae and the sister taxon *S. sichensis*, against the human FUT8 sequence (33,34). Despite the low sequence identity between our algal sequences and FUT8 (∼22% on average), the donor site where the GDP-fucose binds is highly conserved (Figure 8C). This situation was even more apparent when the predicted structure of a representative brown algal protein was superposed on the FUT8 structure, also featuring the hydrolyzed GDP (Figure 8D). The catalytic residues His363, Arg365, Asp368, Lys369 and Ser469, involved in GDP-fucose fixation, are all conserved. This observation indicates that GT23 algal sequences are genuine fucosyltransferases. In contrast the acceptor site shows poor conservation between the algal and human GT23 (Figure 8E). The metazoan GT23s transfer fucose to the innermost GlcNAc residue of N-glycans (34,56). The Src homology 3 (SH3) domain at the C-terminus seems to be involved in protein dimerization, subcellular localization and to some extent, acceptor fixation (57). The SH3 domain is absent from brown algal sequences, and the N-terminal terminus is poorly predicted by AlphaFold2 (and therefore not shown in Figure 8E; the coverage being 89-458 on a total of 458 amino acid sequence). In the *S. latissima* GT23 model, several aromatic residues can be found in close vicinity to the GDP-fucose binding site and might participate in acceptor binding (Figure 8E, yellow residues). Overall, these observations indicate that GT23s in brown algae transfer the fucose residue onto an acceptor molecule distinct from those used by mammalian GT23s. One could speculate that brown algal GT23s catalyse transfer onto oligosaccharidic precursors of FCSPs (*e.g.* FCSOs).

Sulfated fucans are glycans largely restricted to brown algae, exception made of some marine echinoderms (i.e. body wall of sea cucumbers, egg jelly coat of sea urchins) where these molecules are known to be transiently produced and to occur with more linear and regular structures (9,58,59). We therefore also explored the CAZyme content of publicly available echinoderm genomes. We report the presence of the GT10, GT23, GT41 families (Supplementary Table S1). GT10 members are particularly abundant in these organisms, with on average 79 members, compared to 4 GT23 members. The GT23 members in these systems are not very similar to brown algal members of the family and are more closely associated with animal orthologues (Figure 8A). We also screened diatom genomes as recent literature suggest the presence of fucans in these organisms (60). We did not detect any GT23 members in this case and only an average of 3 GT10 members per genome (Supplementary Table S1).

FCSPs, and especially fucans, are known to be heavily sulfated in brown algae. The FCSPs are likely initially polymerized in the state of neutral glycans by GTs, prior to their sulfation by STs. GTs and STs form large multigenic families in brown algae and brown algal STs can be clustered into at least 15 clades (12). In order to investigate the origin of brown algal STs within the Stramenopiles, we analysed the ST gene content of all the Phaeoexplorer brown algal and sister taxa genomes, along with genomes of other Stramenopiles. A phylogenetic tree built using these sequences (Figure 9A, Supplementary Figure S12) confirmed the occurrence of ∼10 major orthologous groups in brown algae. All of these clades could include members with activities on polysaccharides, apart from clade 1, which is expected to be active on non-carbohydrate substrates (7,11,12). Clade 12 includes genes that are located within the syntenic region that also contains the dual FK-GFPP enzymes for the synthesis of the GDP-fucose precursor (Figure 4). This ST and FK-GFPP gene cluster is present in most brown algal genomes, including early diverging ones, suggesting a role for the STs in the sulfation of fucans or fucoidans. In brown algae, clades 4 and 10 include a high number of duplicated genes. Clade 4 includes members from other ochrophyta species but clade 10 appears to be specific to brown algae. In order to study the possible relationship of echinoderm STs with those of brown algae, especially those belonging to clade 10, we clustered these protein sequences using the BLASTP+ Score Ratio (BSR) approach (Figure 9B). The BSR-based heatmap indicates a closer relationship of the brown algal ST sequences from group 10 to echinoderm counterparts than to vertebrate STs, albeit with a pairwise identity score of below 30%. Note that no echinoderm STs have been biochemically characterized so far, thus the conclusion of their possible involvement in fucan sulfation is highly speculative. Regarding the sulfatase genes, as previously reported, brown algae and *S. ischiensis* contain 7 members per genome on average and they all belong to the S1_2 family (18). As for the STs, no biochemical activities have been determined so far for the corresponding enzymes.

**Figure 9.**
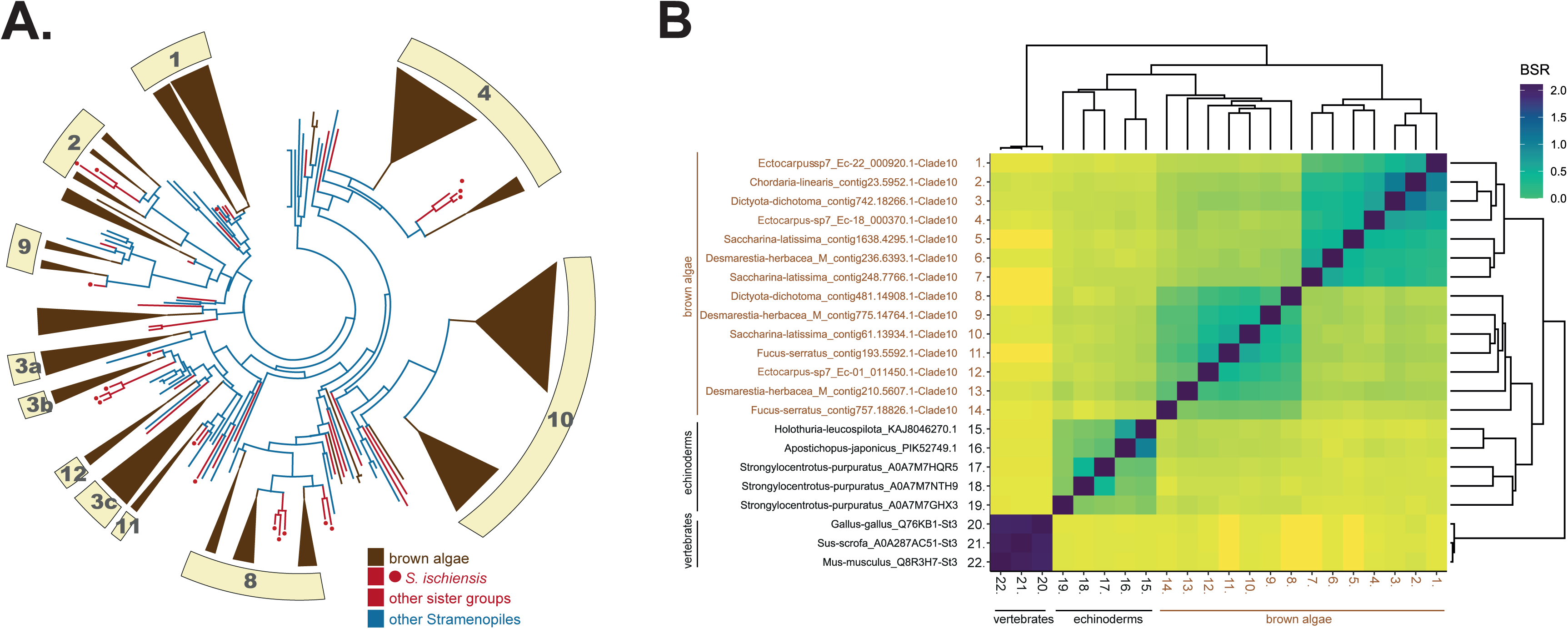
Phylogenetic position of STs from brown algae and other eukaryotes. (**A**) Phylogenetic tree of STs from brown algae and other Stramenopiles. The phylogenetic tree is shown in circle format with collapsed clades. Brown algal sequences are indicated in brown. Sequences belonging to sister groups of brown algae Chrysoparadoxophyceae and Schizocladiaphyceae are indicated by red lines, with red circles for the latter. Other Stramenopile sequences are shown in black. Numbers indicate the brown algal clades as identified in (12). The corresponding uncompressed and fully-labelled tree is shown in Supplementary Figure S12. (**B**) Heatmap representing BLASTP+ Score Ratios (BSRs) for ST protein sequence pairwise alignments. The colour code correlates BSRs. Four groups are apparent: two harbouring brown algal STs (orthologues from group 10), one containing metazoan sequences and the last group containing STs from echinoderms, which is more closely related to brown algal sequences than the metazoan group.

## Discussion

### CAZyme numbers do not correlate with developmental complexity in brown algae

Our analyses provide the first representative view of the core CAZome in brown algae, derived from an exhaustive resource of sequences of distinct taxonomic origins, covering 16 Phaeophyceae families. Prior to the acquisition of large genomic datasets for these organisms, it had been suggested that some CAZyme families, especially GTs, were amplified in algae with more complex tissues (e.g. the kelp *Saccharina japonica*) relative to filamentous forms (e.g. *Ectocarpus* species 7) (61). However, this statement has been gradually invalidated as more genomes have become available (12), and the analysis reported here confirms that the number of total CAZymes in individual brown algal genomes is not correlated with either tissue complexity or taxonomic origin. The filamentous sister taxon *S. ischiensis* only harbors a slightly lower number of CAZymes and the other sister taxa analyzed possessed half as many as *S. ischiensis*. The latter case may indicate the absence of a dense and carbohydrate-rich ECM surrounding these unicellular organisms. One difficulty regarding ‘exotic’ genomes as compared to referenced/model species, lays in the automatic structural annotation of the genes, which may lead to the prediction of fragmented genes. In brown algae, the accounting of domain duplicates in predicted proteins and/or gene fragments by different authors, sometimes in the same organism, can lead to distinct conclusions (12,61,62). One should proceed with caution when drawing conclusions from the analysis of datasets coming from distinct sources. In our specific case, while we cannot address the issue regarding prediction of gene fragments at the whole genome scale, all species have been treated equally from sample preparation to structural and expert annotation, thus limiting the risks in giving biased statements.

Regarding the CAZyme families shared by all brown algae, GT2, GT23, GT31, GT47 and PL41 have a relatively high number of genes per genome, which would be coherent with them being involved in the assembly and remodelling of cell wall polysaccharides. The GT2 family is a large and multispecific family (e.g. includes proteins with different biochemical activities). It is generally one of the largest GT families in eukaryotes, and this is also the case in brown algae. Around two thirds of the GT2 sequences in brown algae are putative cellulose synthase (CESAs) and cellulose synthase-like (CSL) proteins (11), with some being possibly involved in the synthesis of mixed linkage glucans (63). Other GT2s related to dolichyl-phosphate β-glucosyltransferases are involved in protein N-linked glycosylation. The remaining GT2 members do not relate to any specific GT2 homologue with a known activity, and may include putative mannuronan synthases (11). Note that brown algal GT2 genes do not share homology with the Alg8 gene, a GT2 member known to be involved in alginate synthesis in alginate-producing bacteria (11).

GT31 is also a large family in other eukaryotes (*e.g.* 26 genes in *Homo sapiens* and 33 genes in *Arabidopsis thaliana*) with essentially glycoprotein galactosyltransferase and N-acetylglucosaminyltransferase activities and glycolipid activity in metazoans (64), and galactosyltransferase activity during the synthesis of arabinogalactan proteins (AGPs) in plants (31). Similarly, GT47 members are involved in the synthesis of glycosaminoglycans in metazoans and have various activities in the synthesis of cell-wall polysaccharides in plants (32). Brown algae have been shown to express chimeric AGPs (65) and it is tempting to speculate that the GT31 members may be involved in the synthesis of these molecules, although a definite chemical and structural analysis of those glycoproteins still needs to be carried out for brown algae. The PL41 family may be the largest, probably monospecific, CAZyme family in brown algae, indicating the importance of alginate metabolism in these organisms.

### The abilities to synthesize and degrade alginate in brown algae were gained prior to their diversification

Alginate is a major component of brown algal cell walls where its rheological properties plays important roles in determining the wall’s mechanical strength. Our extended analyses confirmed a previous statement (18) that several key gene families involved in alginate synthesis (GMD, ManC5-E) and degradation (PL41, and possibly GH88) were acquired by the common ancestor of brown algae and *S. ischiensis*. This event was followed by a marked amplification of some key families (ManC5-E, PL41) in brown algae relative to its sister taxon.

The ManC5-Es catalyze the final step of alginate synthesis in brown algae: epimerization of mannuronic acid (M) residues into guluronic acid (G) residues, which occurs directly within the polymannuronate polymer. Brown algae are known to contain a large number of ManC5-E genes (12,41,62,66) and this diversity of enzymes presumably yields polymers with a variety of either random or block-wise distribution of G residues, and thus alginates of distinct rheological properties. These modifications are thought to be regulated in response to stress (66), seasonal variations (67) and developmental processes (41). However, this is a one-way process, as the G residues cannot be modified further once generated, thus locking the alginate in a definitive chemical and rheological state. PL41 activity would be relevant in this context, to further modulate the length and the properties of the alginate chains during cell elongation or development. The large size of the PL41 family in brown algae indicates a variety of lytic activities correlated with alginate patterns. As a consequence of these activities, alginates, mannuronan C5-epimerases and alginate lyases, are likely to determine the porosity and mechanical strength of the wall, and may therefore be seen as functional analogues to pectins, pectin methylesterases and pectin lyases in plants, respectively (68).

Alginate is also synthesized by bacteria belonging to the *Azotobacter* and *Pseudomonas* genera (*Gammaproteobacteria*) (69,70). The multigenic mannuronan C5-epimerase family has been extensively studied in these organisms, including at the biochemical level (71). These bacteria also produce alginate lyases, exclusively of the PL5 (72,73) and PL7 families (74,75). *Pseudomonas aeruginosa* is an opportunistic human pathogen, and a leading cause of chronic infection in the lungs of immunocompromised individuals. *Azotobacter vinelandii* is a nitrogen-fixing bacterium found in soil. The ability of these two bacteria to synthesize alginate gives them several selective advantages, the main one being the implementation of a protective shield against adverse environmental conditions (75,76). With the exception of PMM (AlgC), all the genes required for alginate biosynthesis are clustered in a single operon, including PL5-alginate lyase (AlgL) (77,78). The products derived from the lyase action are not used as a carbon source by the bacteria, thus several functions have been putatively attributed to these lyases (76,77). In *P. aeruginosa* they contribute to cell viability by clearing the periplasmic space of trapped alginate (76,78). In *A. vinelandii* they trigger coat rupture during cyst germination (74). Additional alginate lyase activities have been described in other PL families, essentially from bacteria (PL6, PL15, PL17, PL18, PL31, PL32, PL36, PL39) and some eukaryotes (PL8, PL14) (12). Screening of public databases indicates uncharacterized PL41 homologues in bacteria, essentially from the *Streptomyces* within the *Actinomycetota* (Figure3A, Supplementary Figure S4) but these organisms are not known to contain an alginate biosynthetic pathway. Indeed, in contrast to the *Azotobacter* and *Pseudomonas* genera, most of these bacteria use the products of alginate lyase action as a carbon source (79).

The occurrence of alginate metabolic pathways in *S. ischiensis* correlates with the immunodetection of alginate in this species (19). There is strong evidence that the ManC5-E, PL41 and GH88 genes have been acquired by *S. ischiensis* via horizontal gene transfer (HGT). We have previously proposed that about 1% of brown algal genes are derived from HGTs, with these genes having been principally acquired from bacterial genomes (18). Around 10% of those HGT-derived genes have predicted functions in carbohydrate transport and metabolism (18).

While previous work indicated that brown algal GMD was derived from a HGT of an ancestral GMD sequence from *Actinobacteria* (11), our analyses based on additional algal sequences indicates that this conclusion is not strongly supported, and other origins of horizontal gene transfer cannot be excluded, in particular from other bacterial groups. Surprisingly, the brown algal ManC5-E and PL41 families seem to derived from distinct HGTs (different donor taxa). Thus, the catalytic steps for the synthesis and the degradation of alginate, may have arisen independently in *S. ischiensis* and brown algae, although within a narrow time-scale. While a global CAZome expansion is not observed in brown algae as compared to *S. ischiensis*, the PL41 family is the exception to this rule, with a greater number observed in brown algae as compared to its sister taxon (i.e. an average of 22 genes vs. 3). The situation is similar for the ManC5-E family which shows great expansion in brown algae (i.e. an average of 27 genes vs. 11). The acquisition of these enzymatic steps, and the expansion of the gene families in brown algae, probably allowed the generation of extensive ECMs with fine-tuned structures and rigidity. While there is no dynamic correlation between the number of CAZyme families and tissue complexity, these extensive ECMs may have represented an important prerequisite for the evolution of developmental complexity, and for the emergence of large, resilient substrate-anchored multicellular organisms, in the hostile coastal environment.

### The fucan biosynthetic pathway in brown algae needs to be fully resolved

The set of genes involved in the synthesis of sulfated glycans remains unknown. Access to the large Phaeoexplorer genome dataset allowed us to identify the most promising candidate genes. Based on the large size of the GT23 family in all brown algal genomes, the specific phylogenetic position of the brown algal members of this family compared to biochemically characterized members, and information gained from modelling of GT23 protein structure, this family represents a strong candidate for fucan synthases in brown algae. This hypothesis is also supported by transcriptomic data obtained during the development of the *Fucus* embryo (80), where the expression of GT23 genes is up-regulated during early fucan deposition into the walls. Elongation of the oligosaccharidic moieties of FCSPs is believed to be followed by sulfation, catalysed by STs. The diversity of the brown algal ST orthologue groups may indicate a variety of acceptors, including many monosaccharides and their available hydroxyl positions found in FCSPs. In contrast to the GTs, we found homologues of some brown algal STs in echinoderms and these enzymes are known to synthesize sulfated fucans of regular patterns (58). While the origin of such homology might find its roots near the origin of eukaryotes (LECA), this would imply that the ability to perform fucan elongation and its subsequent sulfation, to have been lost in all eukaryotes, except brown algae and a narrow number of taxa within animals. The current available data does not identify taxa donors to favour a scenario involving HGT in the two groups of organisms. Functional convergence could also explain part of the similarity, yet this needs to be biochemically explored. STs belonging to orthologue group 10 are interesting candidates for the sulfation of fucans but other groups cannot be excluded. For example, orthogroup 12 contains a gene that is frequently associated with FK-GFFP in a syntenic region, and groups 2 and 8 contain genes that are also up-regulated during early development of the *Fucus* embryo (80). These latter two groups contain orthologues from *S. ischiensis*. At this stage we cannot conclude that the GT23 family, the STs, the SAs and the fucan biosynthetic pathway were obtained independently in brown algae as compared to echinoderms. Functional validation of all of those genes is needed to progress further on this question.

In addition to brown algae and echinoderms, immunodetection procedures and fucose assays have provided evidence for the presence of fucans/fucoidans in the diatoms *Chaetoceros* and *Thalassiosira weissflogii* (81). The composition of these secreted glycans in diatoms differ markedly from those of brown algal fucans/fucoidans (82). The precise chemical structure of the diatom glycans has not been investigated in detail, the structures reported being essentially limited to glucuronomannans (83). It is therefore possible that the anti-fucans/fucoidans antibodies cross-react with additional, yet to be described, glycan structures in these organisms. The BAM2 antibody, which recognises α-(1→3)-fucans featuring 4-O-sulfate esters based on analysis using synthetic oligofucans (60), detects epitopes in diatoms, but precise structural validation of the compounds detected still needs to be carried out. In addition, glycan content has not been explored in detail in Stramenopiles other than brown algae and diatoms. A few studies have reported analyses of total monosaccharide composition in some of these organisms and these analyses either did not detect fucose (*Tribonema minus,* Xanthophyceae) (84), or only detected fucose at a limited level (*Nannochloropsis oceanica* Eustigmatophyceae) (85). These results suggest that FCSPs are unlikely to be constitutively produced by these organisms at high levels. In short, both the lack of biochemical validations of any putative fucan synthase genes, and the fragmented knowledge of FCSP structures outside brown algae, impairs progression on tracking the evolutionary history of fucan synthesis in eukaryotes.

The situation of the sister taxon *S. ischiensis* is intriguing. Alginate has been detected immunologically in this species (19) and all the genes involved in alginate biosynthesis and degradation in brown algae are conserved. We have also identified GT23 members highly similar to their brown algal orthologues (∼48% protein identity). The ability of *S. ischiensis* to produce sulfated fucans needs to be established. The outcome would contribute towards testing the hypothesis that the GT23 family contains fucan synthase genes. If future research establishes the occurrence of fucans in *S. ischiensis*, this would indicate an ECM with a glycan composition very similar to that of brown algae. One substantial difference between the Phaeophyceae and the Schizocladiophyceae is the absence of plasmodesmata in *S. ischiensis* (19,86). Plasmodesmata have been shown to be crucial in cell-to-cell signaling in land plants and may play a role in the evolution of tissue complexity (87–89). Thus, it is not only the building up of a consistent and dynamic glycan-rich ECM in brown algae that may have contributed to the evolution of complex multicellular organisms, but also some structural features that function in close vicinity to ECMs such as plasmodesmata.

## Acknowledgements

We are grateful to the Roscoff Bioinformatics platform ABiMS (http://abims.sb-roscoff.fr), part of the Institut Français de Bioinformatique (ANR-11-INBS-0013) and BioGenouest network, for providing computing and storage resources.

## Author contributions

Software: L.B.G., E.C.; Formal analysis: F.D., O.G. and J.M.C; Investigation: L.M., N.T., A. L. and C.H; Resources: E.C. , J.M.C. and N.T.; Data Curation: T.B. and N.T.; Writing-Original Draft: L.M., A. B.-C. and C.H.; Writing-Review & Editing: all authors; Visualization: L.M., A.B-C., M.C., A.L. and C.H.; Supervision: C.H.; Project administration: F.D., J.M.C. and C.H.; Funding acquisition: J.M.C. and C.H.; Conceptualization: C.H.

## Funding

This work was supported by the ANR project BrownSugar (ANR-20-CE44-0011) and the France Génomique National infrastructure project Phaeoexplorer (ANR-10-INBS-09).

## Supplementary Figure legends

**Supplementary Figure S1.** Phylogenetic trees of key enzymes involved in the synthesis of GDP-mannose, a precursor for the synthesis of both alginates and fucans. (**A**) The bifunctional annose-6-phosphate isomerases (MPI)/mannose-1-phosphate guyanylyltransferases (MPG) and (**B**) the bifunctional phosphomannomutases (PMM)/ phosphoglucomutase (PGM).

**Supplementary Figure S2.** Clustering and phylogenetic position of brown algal proteins within the GDP-mannose/UDP-glucose dehydrogenases (GMD/UGDs) superfamily. (**A**) Heatmap representing BLASTP+ Score Ratios (BSRs) for protein sequence pairwise alignments. The colour code correlates BSRs. Two main groups of Stramenopile sequences are clearly apparent: one containing the GMD sequences and one harbouring the UGD sequences. The bacterial sequences from *Actinobacteria* and *Pseudomonadota* are distantly related to these two groups. (**B**) Uncompressed version of the phylogenetic tree of GMDs shown in Figure 2A of the main manuscript.

**Supplementary Figure S3.** Uncompressed version of the phylogenetic tree of ManC5-E proteins shown as an inset in Figure 2B of the main manuscript

**Supplementary Figure S4.** Uncompressed version of the phylogenetic tree of PL41 proteins shown in Figure 3A of the main manuscript

**Supplementary Figure S5.** Phylogenetic tree of DEHU reductases, known to be active in alginate degradation in brown algae

**Supplementary Figure S6.** Uncompressed versions of the phylogenetic tree of GH88 proteins shown in Figure 3B of the main manuscript

**Supplementary Figure S7.** Phylogenetic trees of key enzymes involved in the synthesis of GDP-fucose. Enzymes involved in the synthesis of GDP-fucose through the *de novo* pathway: (**A**) GDP-mannose 4,6-dehydratases (GM46D) and (**B**) the GDP-L-fucose synthetases (GFS); and through the salvage pathway: (**C**) fucokinase (FK) and (**D**) GDP-fucose pyrophosphorylase (GFPP).

**Supplementary Figure S8.** Uncompressed version of the phylogenetic tree of GT10 proteins shown in Figure 5A of the main manuscript

**Supplementary Figure S9.** Uncompressed version of the phylogenetic tree of GT41 proteins shown in Figure 6 of the main manuscript

**Supplementary Figure S10.** Uncompressed version of the phylogenetic tree of GT74 proteins shown in Figure 7 of the main manuscript

**Supplementary Figure S11.** Uncompressed version of the phylogenetic tree of GT23 proteins shown in Figure 8B of the main manuscript

**Supplementary Figure S12.** Uncompressed version of the phylogenetic tree of ST proteins shown in Figure 9A of the main manuscript.

**Supplementary Table 1.** Gene number of a selection of GT families in brown algae and other relevant organisms and known to contain FucT activities in the CAZy database. All known FucTs are classified in the CAZy GT families as shown. The generated glycosidic linkages identified from characterized enzymes (outside brown algae) are indicated. The average value is shown for the following species: *Ectocarpus* sp.7, *Pylaiella littoralis, Chordaria linearis* (Ectocarpales), *Saccharina latissima, Saccharina japonica, Macrocystis pyrifera* (Laminariales), *Porterinema fluviatiles* (Ralfsiales), *Ascophyllum nodosum, Fucus serratus, Fucus distichus* (Fucales), *Saccorhiza dermatodea, Saccorhiza polyschides* (Tilopteridiales), *Desmarestia herbacea*, *Desmarestia dudresnayi* (Desmarestiales), *Sphacelaria rigidula* (Sphacelariales), *Dictyota dichotoma* (Dictyotales), *Discosporangium mesarthrocarpum, Choristocarpus tenellus* (Discosporangiales), *Schizocladia ischiensis* (Schizocladiophyceae), *Heterosigma akashiwo* (Raphidophyceae), *Phaeodactylum tricornutum CCAP 1055, Thalassiosira pseudonana* CCMP 1335*, Chaetoceros tenuissimus* NIES-3715_426638 (diatoms), *Lytechinus variegatus* NC3_7654, *Strongylocentrotus purpuratus* Spur-01_7668 (sea urchin), *Apostichopus japon* Shaxun_307972 (sea cucumber). ND. Not determined.

| GT families | GT10 | GT23 | GT41 | GT74 | GT11 | GT37 | GT65 | GT68 |
| --- | --- | --- | --- | --- | --- | --- | --- | --- |
| Known FucT activities in CAZy | α-1,3/4-FucTs | α-1,6-FucTs | protein O-FucTs | α-1,2-FucTs | α-1,2/3-FucTs | α-1,2-FucTs | protein O-FucTs | protein O-FucTs |
| **Phaeophyceae** |  |  |  |  |  |  |  |  |
| Ectocarpales [3] | 1 | 9 | 13 | 1 | 0 | 0 | 0 | 0 |
| Laminariales [3] | 1 | 9 | 10 | 1 | 0 | 0 | 0 | 0 |
| Ralfsiales [1] | 3 | 3 | 12 | 1 | 0 | 0 | 0 | 0 |
| Fucales [3] | 1 | 14 | 10 | 1 | 0 | 0 | 0 | 0 |
| Tilopteridiales [2] | 0 | 5 | 10 | 2 | 0 | 1 | 0 | 0 |
| Desmarestiales [2] | 1 | 11 | 16 | 1 | 0 | 1 | 0 | 0 |
| Sphacelariales [1] | 0 | 1 | 6 | 1 | 0 | 0 | 0 | 0 |
| Dictyotales [1] | 1 | 4 | 10 | 1 | 0 | 1 | 0 | 0 |
| Discosporangiales [2] | 1 | 1 | 5 | 1 | 0 | 1 | 0 | 0 |
| **Sister groups to Phaeophyceae** |  |  |  |  |  |  |  |  |
| Schizocladiphyceae [1] | 1 | 5 | 11 | 1 | 0 | 1 | 0 | 0 |
| Raphidophyceae [1] | 0 | 0 | 8 | 0 | 0 | 0 | 0 | 0 |
| **Other Stramenopiles** |  |  |  |  |  |  |  |  |
| diatoms [3] | 3 | 0 | 2 | 1 | ND | ND | ND | ND |
| **Echinoderms** |  |  |  |  |  |  |  |  |
| Sea urchin [2] | 60 | 7 | 4 | 0 | ND | ND | ND | ND |
| Sea cucumber [1] | 97 | 1 | 1 | 0 | ND | ND | ND | ND |

## Notes

### Competing Interest Statement

The authors have declared no competing interest.

### Summary of Updates

This includes only minor revisions. The abstract has been shorten and some layouts have been updated.

