## Supplementary Figures for "Evolutionary history of the main extracellular matrix polysaccharides in brown algae"

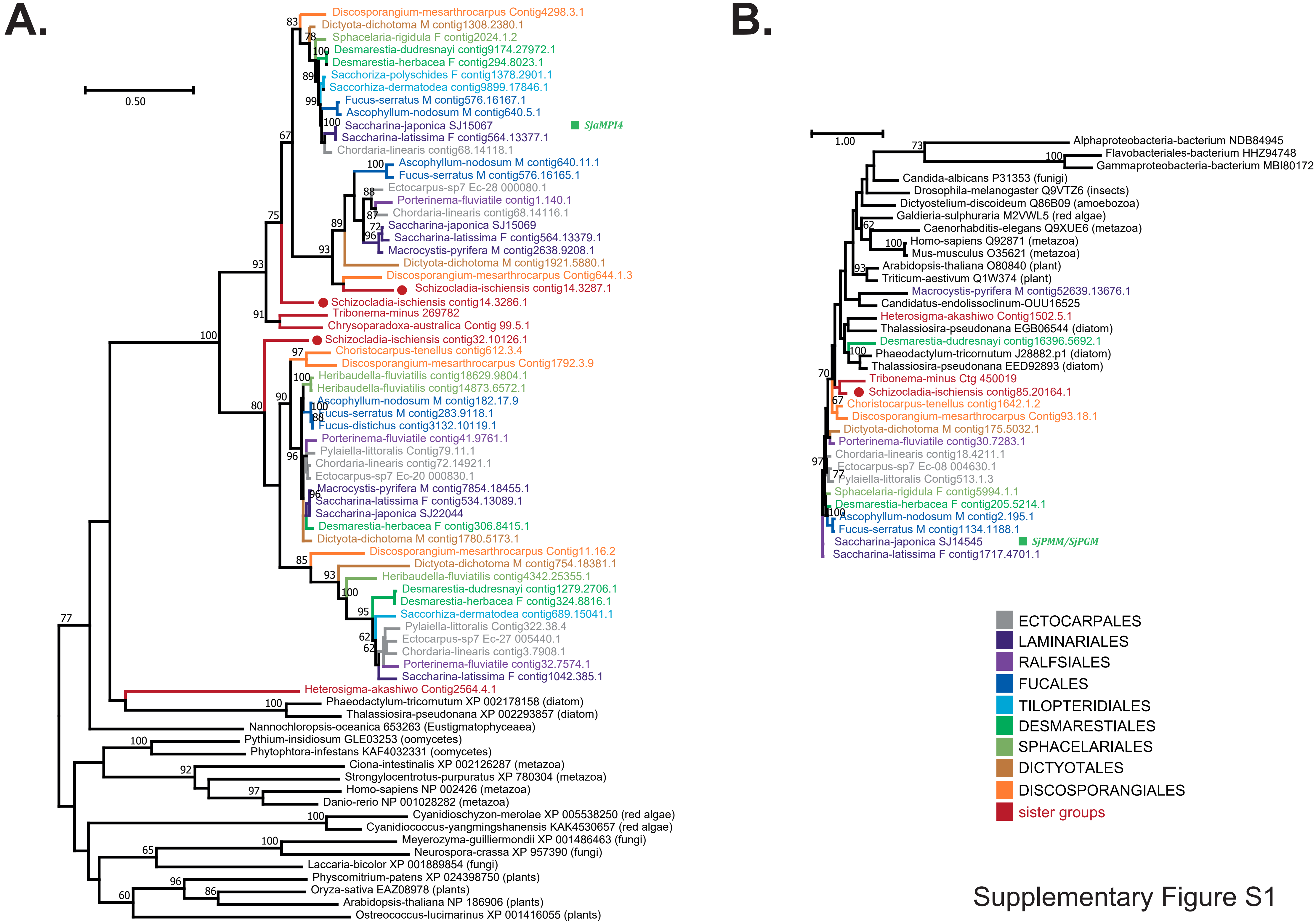


Supplementary Figure S1

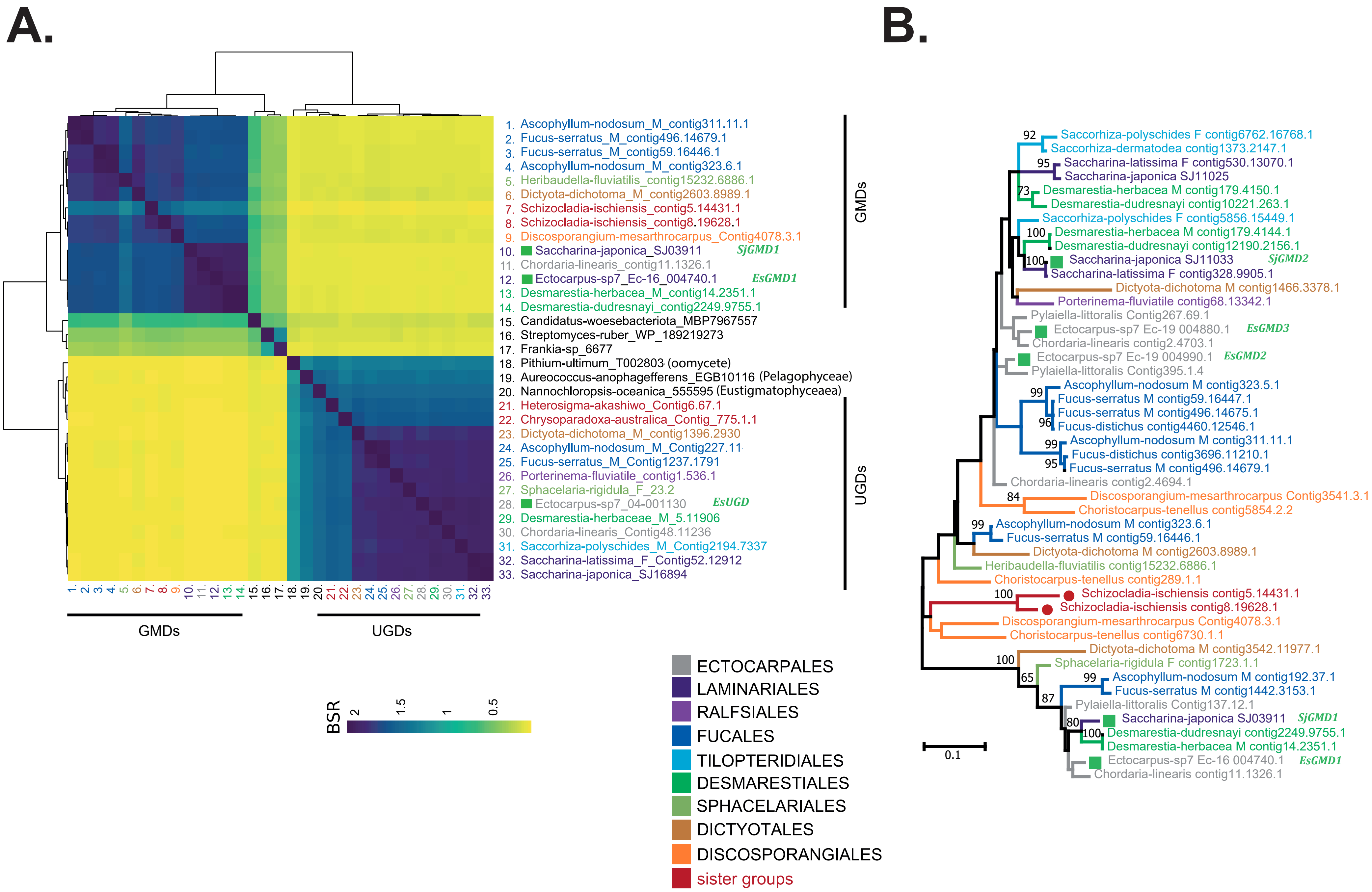


Supplementary Figure S2

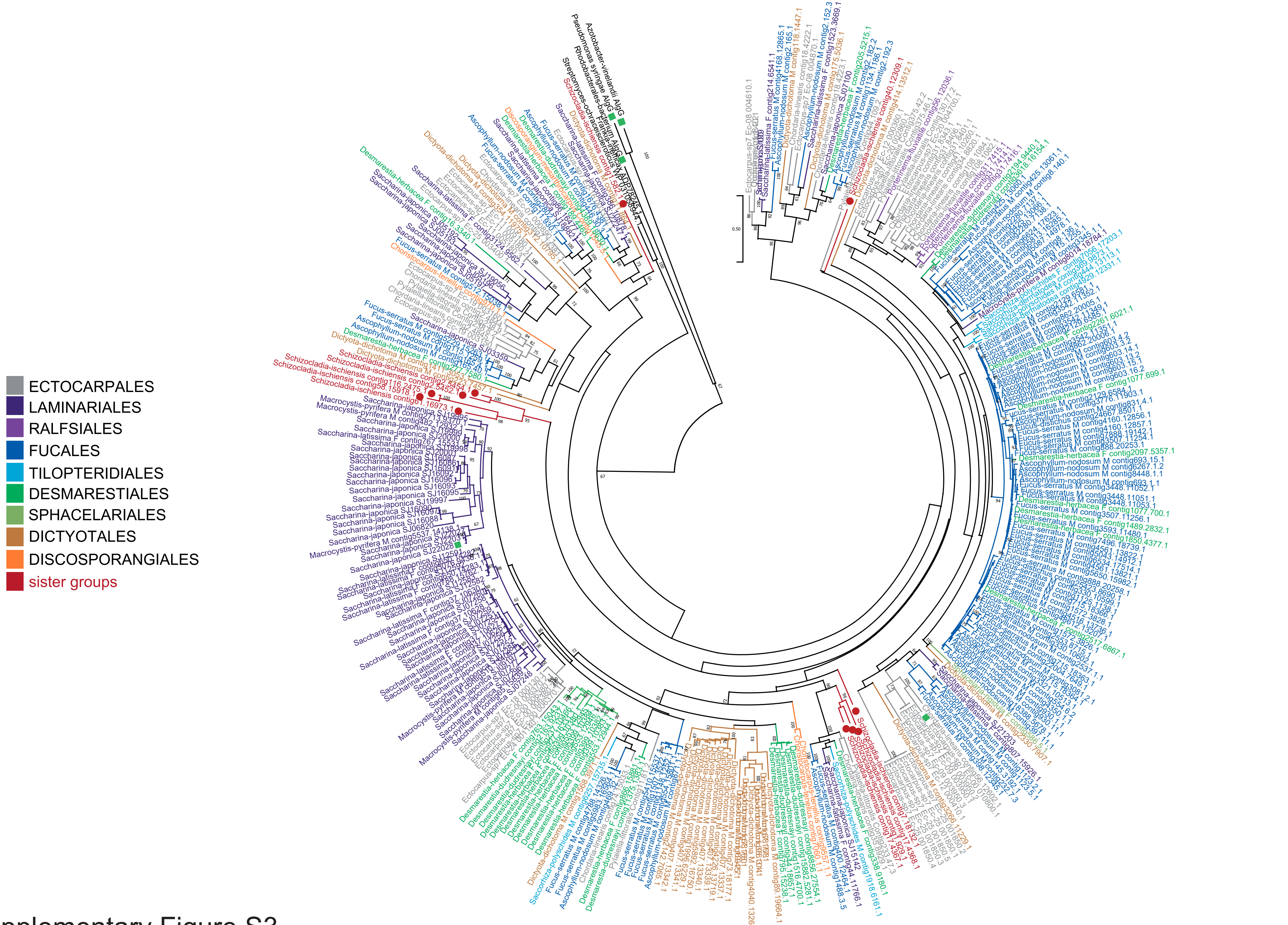


Supplementary Figure S3

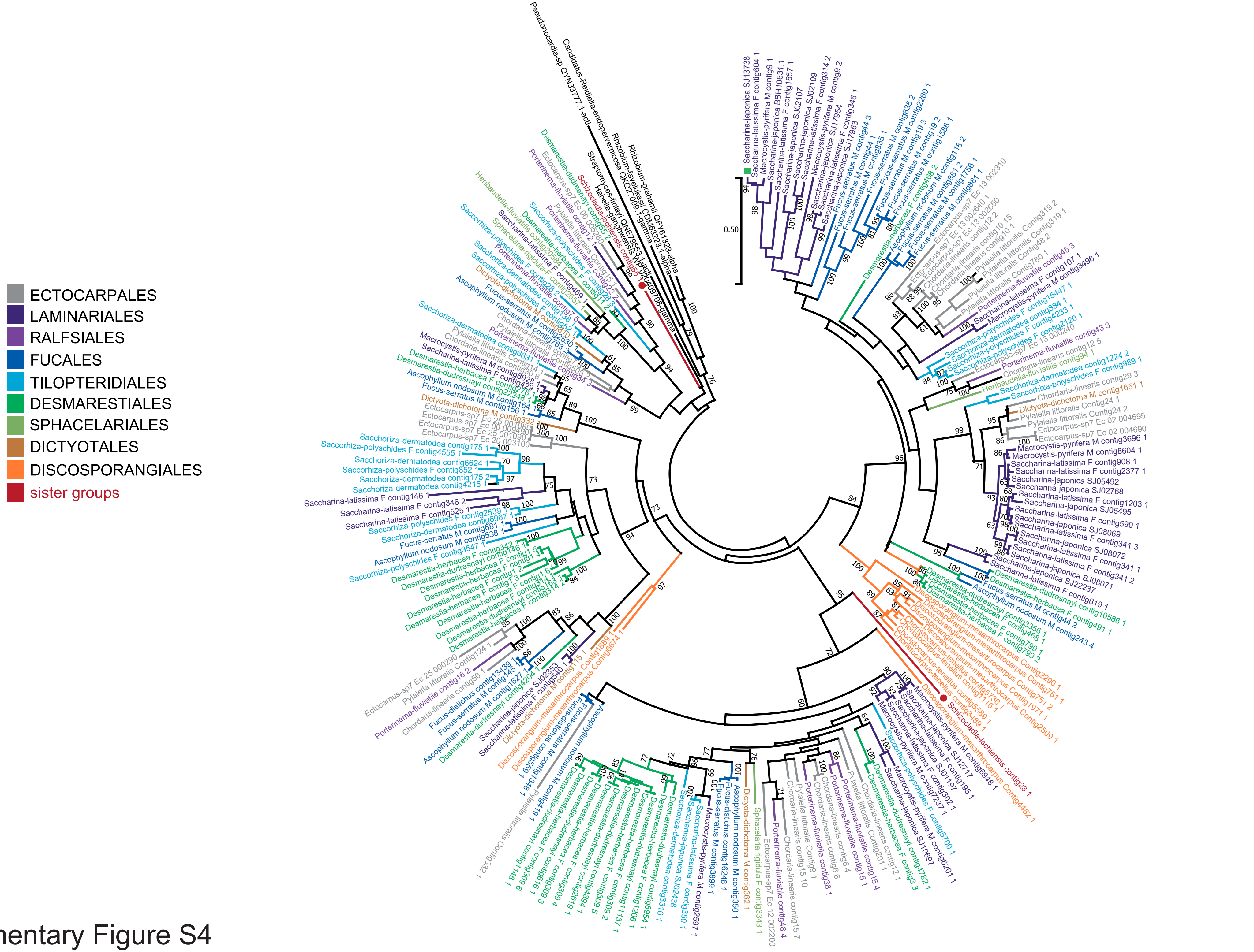


Supplementary Figure S4

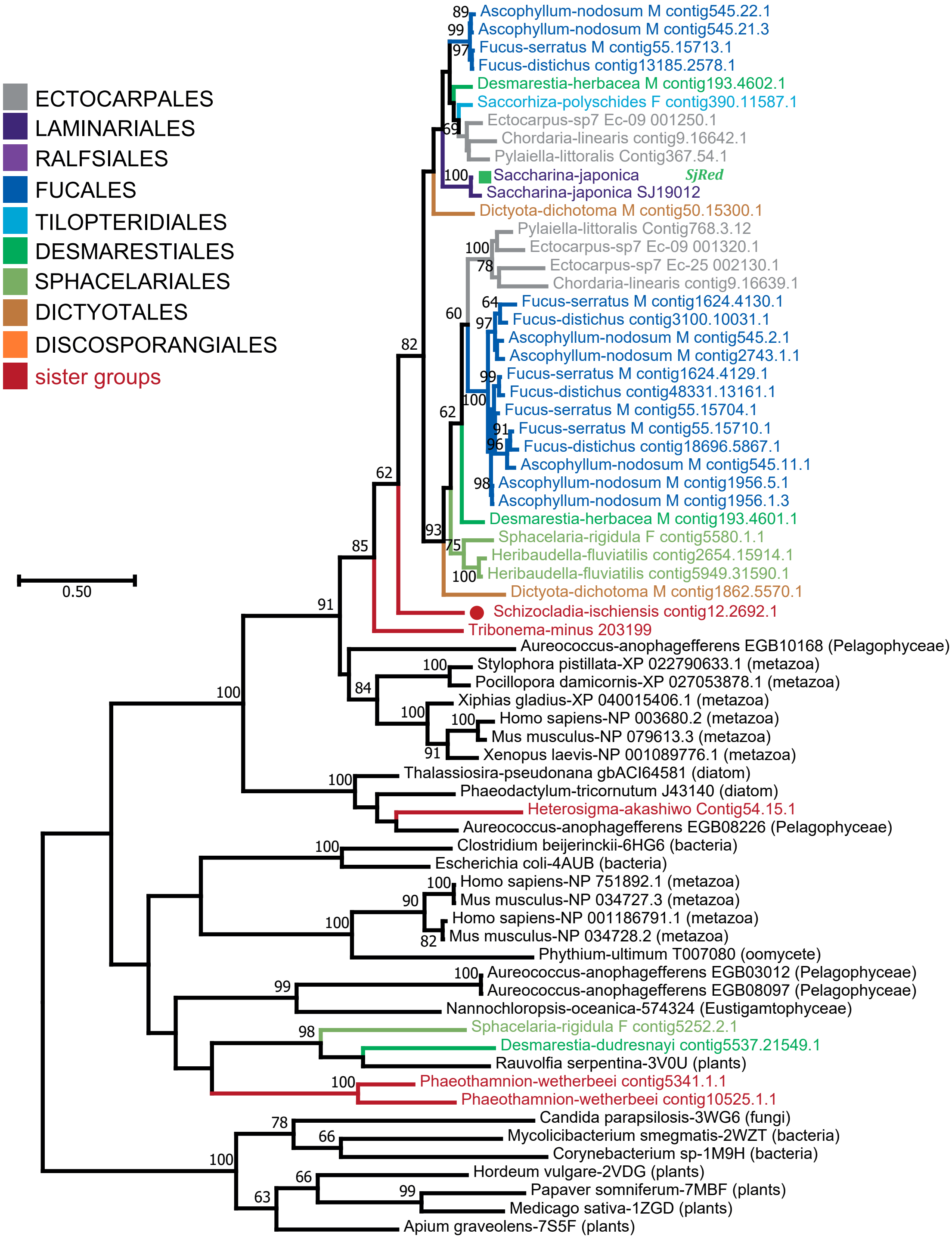


Supplementary Figure S5

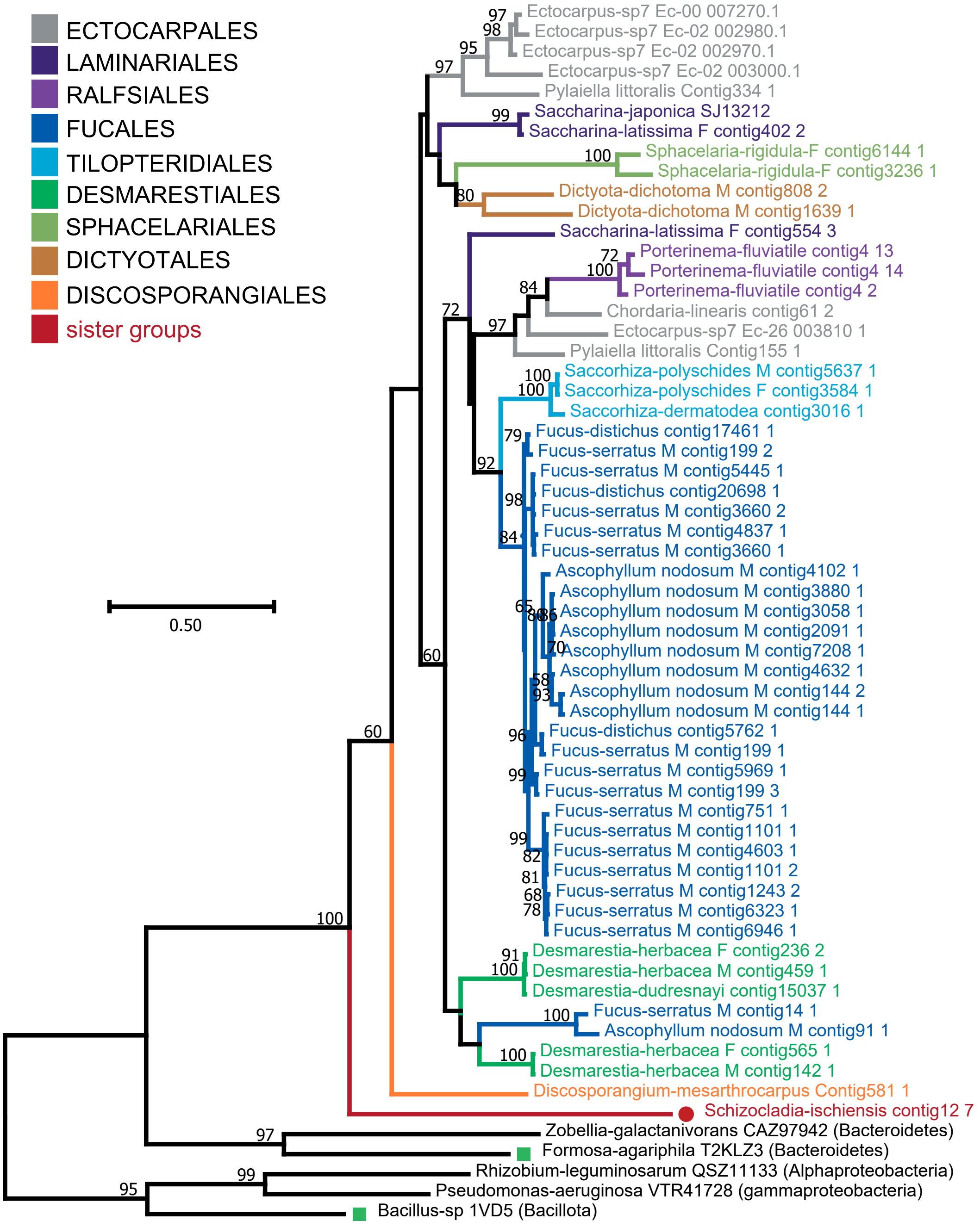


Supplementary Figure S6

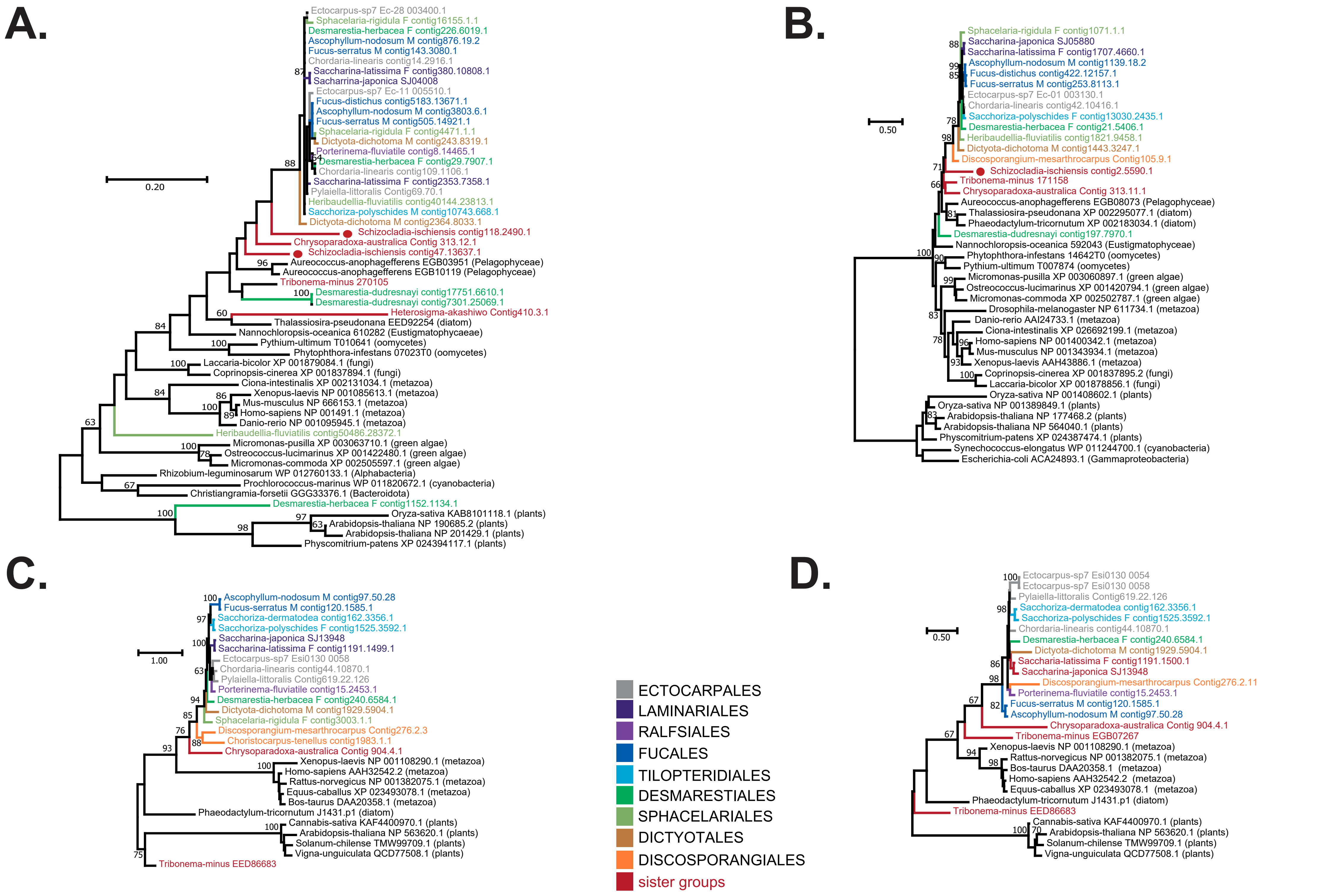


Supplementary Figure S7

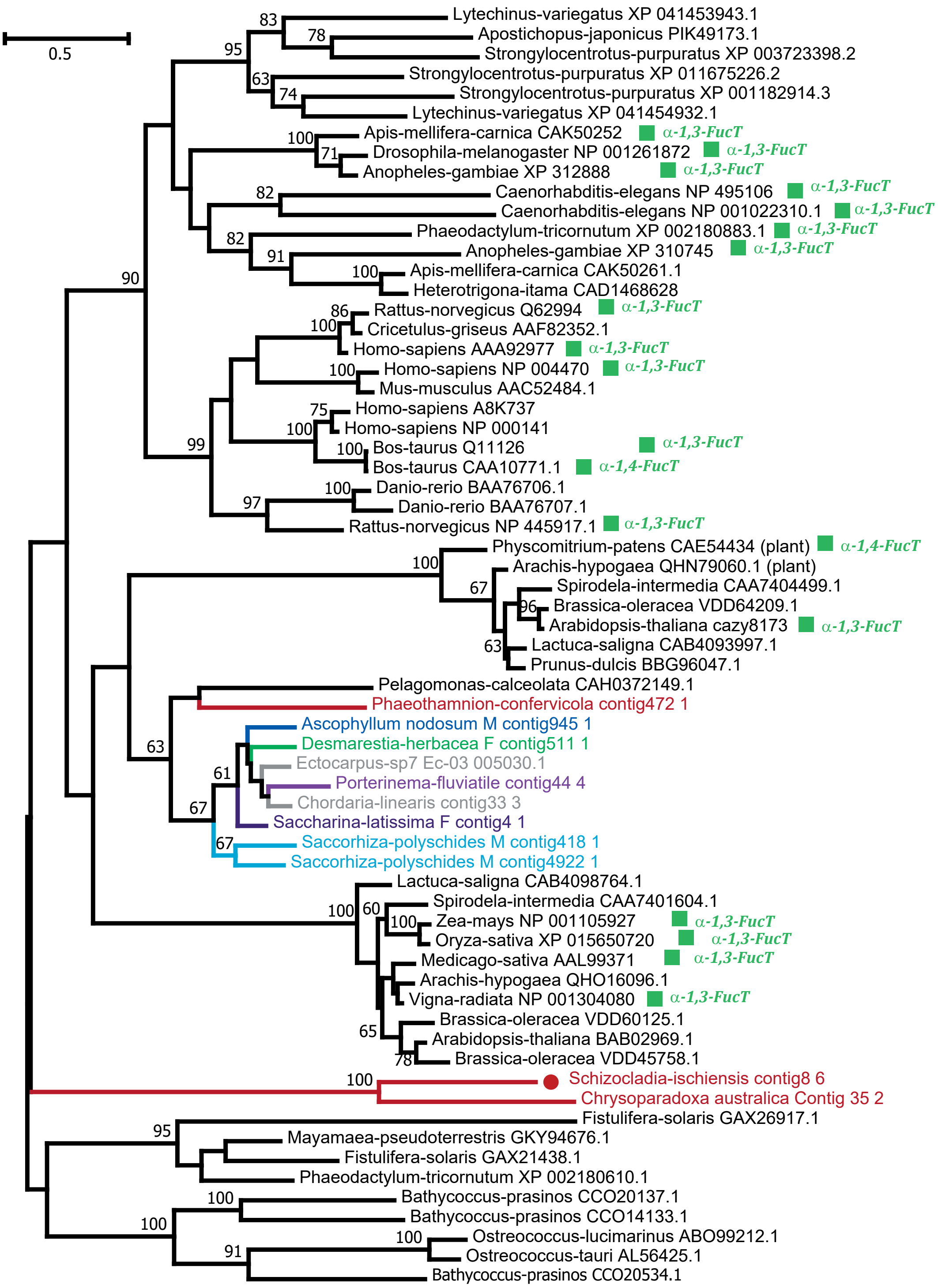


Supplementary Figure S8

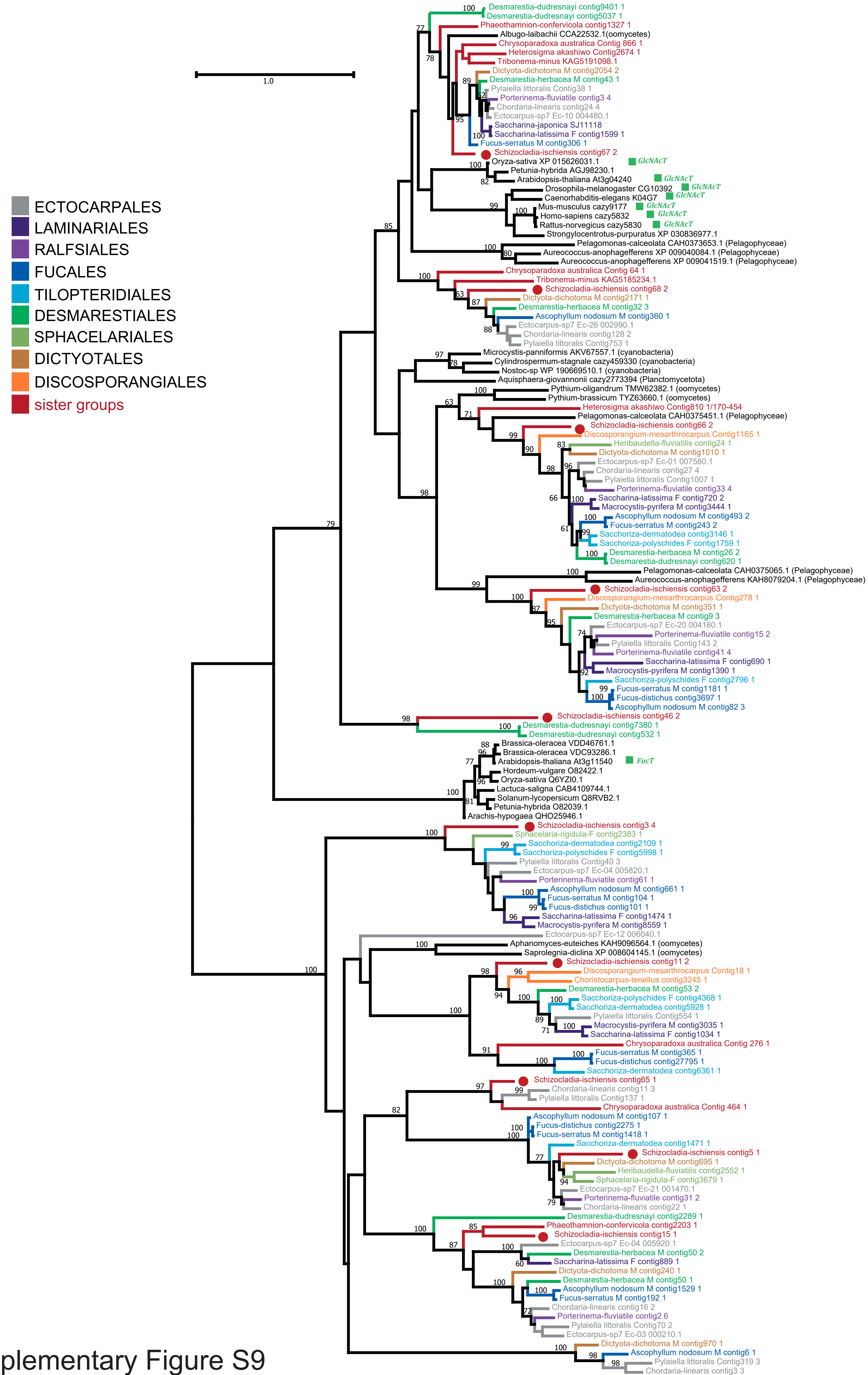


Supplementary Figure S9

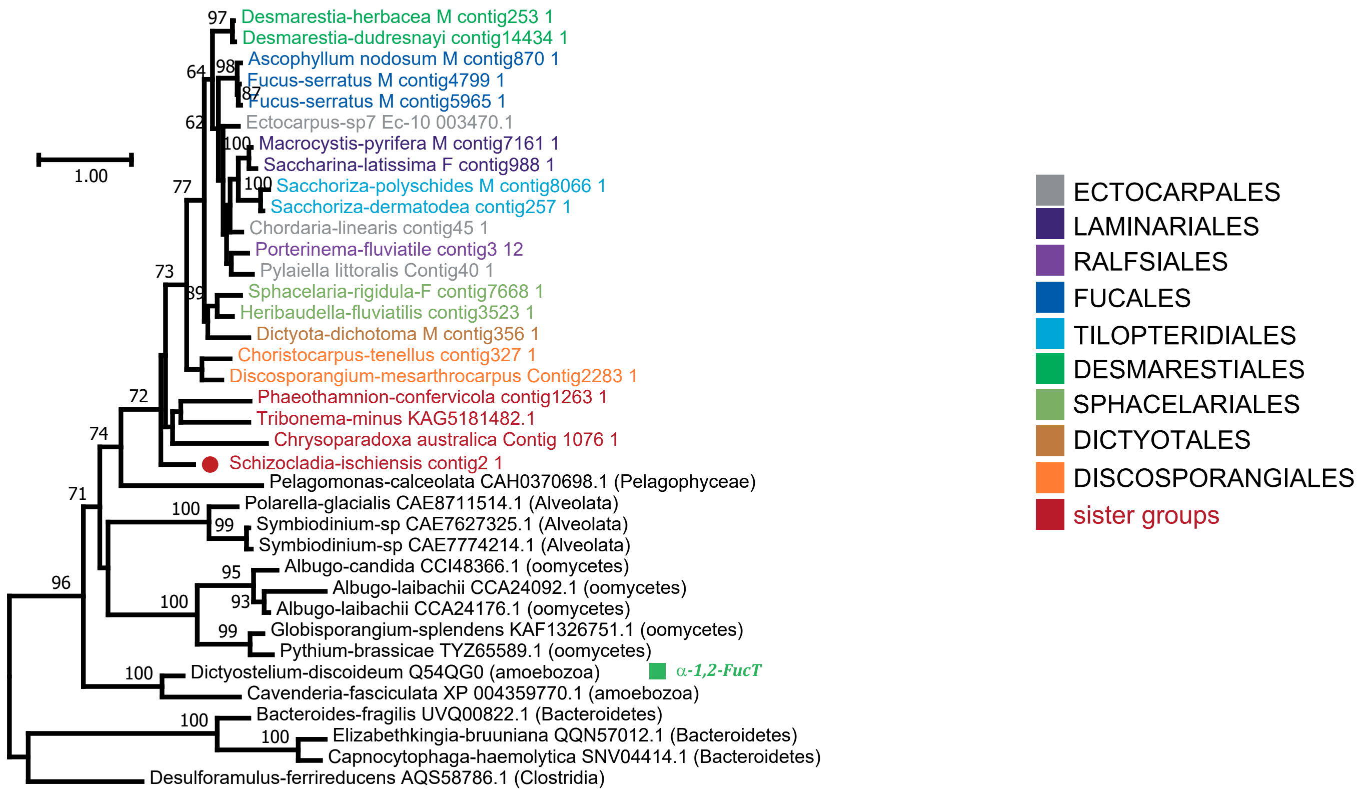


Supplementary Figure S10

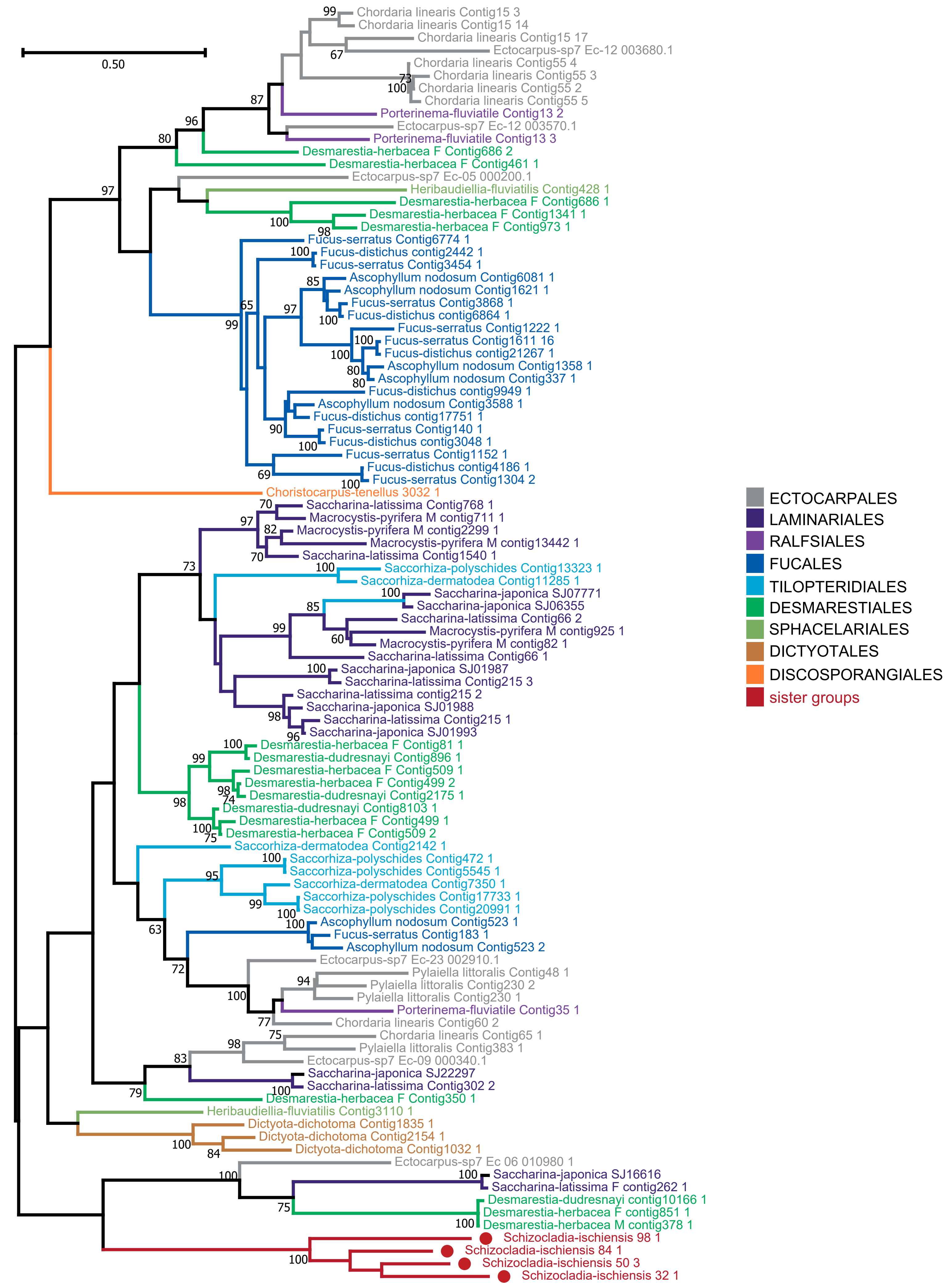


Supplementary Figure S11

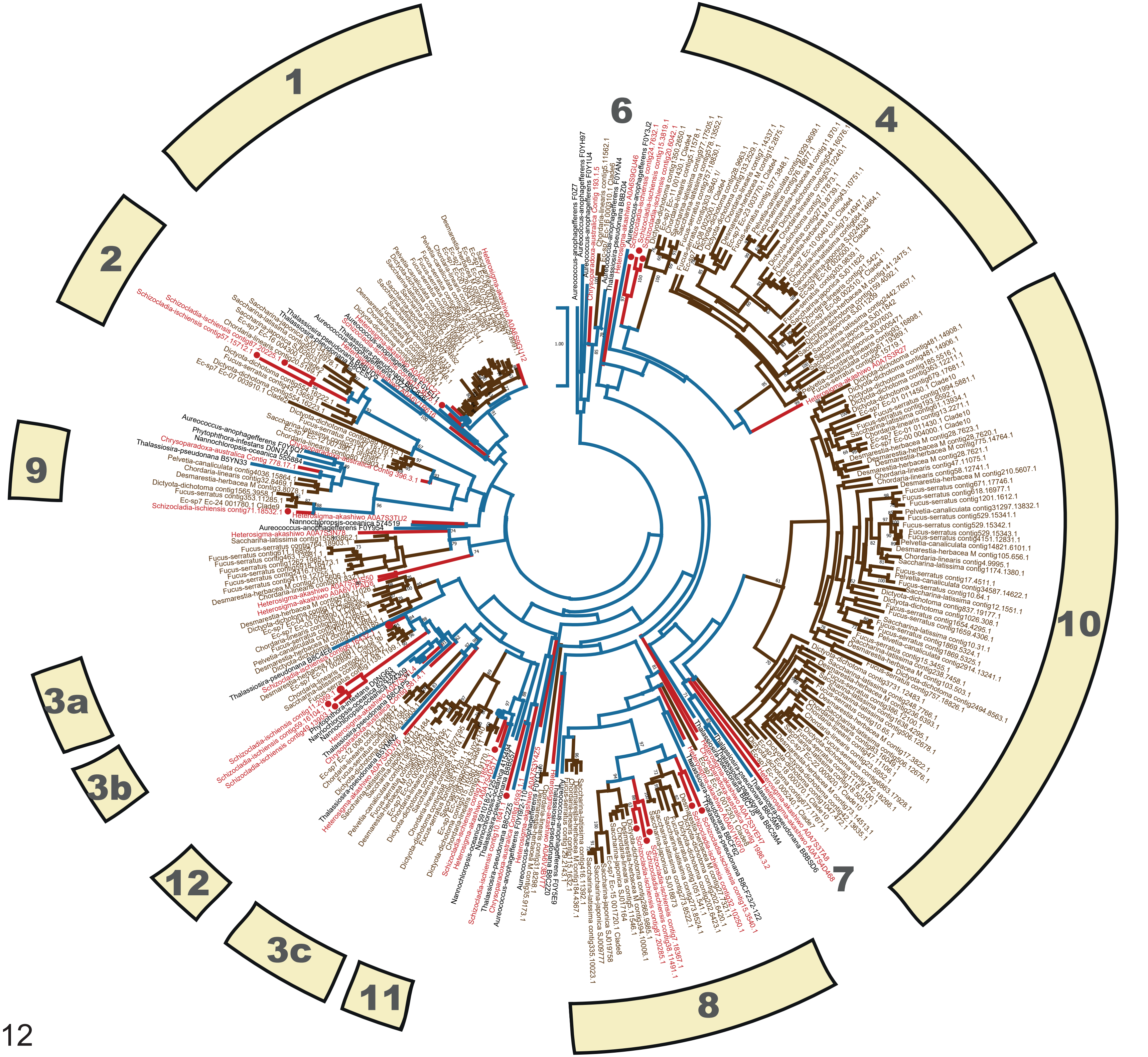


Supplementary Figure S12
